# The β-lactam adjuvant guanosine potentiates anti-folate antibiotics and pyrimidine synthesis inhibitors by depleting thymidine in methicillin-resistant *Staphylococcus aureus*

**DOI:** 10.64898/2026.01.15.699642

**Authors:** Aaron C. Nolan, Jessica B. Kelly, Jongsam Ahn, Dhananjay Shinde, Vinai C. Thomas, Merve S. Zeden, James P. O’Gara

## Abstract

Efforts to improve the effectiveness of existing interventions for antimicrobial-resistant (AMR) infections include identifying new ways to overcome resistance to licensed antibiotics using adjuvants or deploying antibiotics in novel combinations. Although antibiotics targeting the bacterial cell wall (e.g., β-lactams) and folate metabolism (e.g., trimethoprim-sulfamethoxazole, TMP-SMX) remain cornerstones of modern healthcare, resistance to both classes poses an ongoing therapeutic challenge. We recently demonstrated that purine nucleosides can act as potent antibiotic adjuvants, restoring β-lactam susceptibility in methicillin-resistant *Staphylococcus aureus* (MRSA). Here, we show that the β-lactam adjuvant guanosine significantly reduces intracellular thymidine levels in MRSA and potentiates the activity of antifolate antibiotics (TMP-SMX) as well as the pyrimidine antimetabolites 5-fluorouracil (5-FU) and 5-fluorouridine (5-FUrd). Incorporation of oxacillin into guanosine-antifolate or guanosine-pyrimidine analogue combinations further enhanced killing of both planktonic and biofilm-associated MRSA. Thymidine depletion was accompanied by elevated intracellular reactive oxygen species (ROS) and dissipation of membrane potential, providing mechanistic insight into the bactericidal effects of these combinations. Together, these findings demonstrate that guanosine expands MRSA susceptibility beyond β-lactams to include additional clinically relevant antimicrobial drug classes commonly used to treat bacterial infections.

**Author Summary.:** Finding new ways to help antibiotics to work against resistant bacteria is an important part of efforts to address the antimicrobial resistance crisis. Adjuvants, which are boosters that help antibiotics to work against resistant bacteria, are important in this regard. We recently discovered that, guanosine, which is involved in the synthesis of DNA, makes methicillin-resistant *Staphylococcus aureus* (MRSA) more sensitive to penicillin-type antibiotics. Here we report that guanosine also improves the effectiveness of two other types of antimicrobial drugs (anti-folate antibiotics and pyrimidine antimetabolites) that are also used in clinical practice, expanding the potential of this adjuvant to improve the management of difficult-to-treat MRSA infections.

## Introduction

Finding new ways to preserve the efficacy of currently-licensed antimicrobial drugs is a central part of efforts to address the antimicrobial resistance (AMR) crisis. In addition to new drug discovery, which is costly and time-consuming, repurposing currently-licensed antibiotics in new combinations and the discovery of novel adjuvants also have significant potential to improve treatment options for AMR infections. Antibiotic adjuvants, which potentiate antibiotic activity or disrupt resistance mechanisms, have proven clinical usefulness. In 2023 we reported that the purine nucleosides guanosine (Gua) and xanthosine (Xan) have activity as adjuvants that can resensitise methicillin resistant *Staphylococcus aureus* (MRSA) to oxacillin (OX) and other β-lactam antibiotics (1). Exposure of MRSA to Gua significantly reduced the levels of the cyclic dinucleotide c-di-AMP, which is required for β-lactam resistance (1). In contrast exposure of MRSA to adenosine (Ade) did not significantly effect c-di-AMP levels and instead increased OX resistance. Inosine, which can be fluxed into the ATP or GTP branches of purine metabolism, only marginally affected OX resistance (1). c-di-AMP controls osmotic regulation and exposure of MRSA to OX/Gua was accompanied by a 2-fold increase in cell size (1). These data support the conclusion that perturbation of purine homeostasis by exogenous nucleosides controls MRSA resistance to β-lactams. Drugs derived from nucleotides are used in the treatment of cancer and viral infections raising the possibility that purine nucleosides may potentiate the activity of β-lactams against MRSA.

In this work, we compared the impact of exposure to OX/Gua or OX/Ade combinations on the MRSA metabolome and revealed significant pleiotropic effects on cell wall, purine and pyrimidine (specifically thymidine) metabolism. Thymidine biosynthesis is dependent on tetrahydrofolate (THF), which serves as a co-factor for thymidylate synthase (ThyA)-mediated conversion of uracil into thymidine (2–4), suggesting that purine adjuvants may potentiate the activity of the anti-folate antibiotics trimethoprim-sulfamethoxazole (TMP-SMX), as well as the pyrimidine analogue anti-cancer drugs 5-fluorouracil (5-FU) and 5-fluorouridine (5-FUrd). TMP-SMX, which are used in a 1:5 ratio combination, block the activity of dihydrofolate reductase and dihydropteroate synthase respectively, required for THF synthesis. This formulation also known as bactrim and co-trimoxazole is commonly used to treat MRSA (5) and several other bacterial infections (6). However, resistance is observed in the clinic, especially in chronic *S. aureus* infections with the rise of thymidine-dependent small colony variants, with mutations in *thyA* (2–4, 7, 8).

Reduced thymidine levels in OX/Gua-treated MRSA raised the possibility that purine adjuvants may also potentiate the activity of antibiotics that interfere with thymidine and/or pyrimidine biosynthesis. To investigate this, synergy between Gua and the anti-folate antibiotics sulfamethoxazole (SMX), trimethoprim (TMP), and the pyrimidine analogues 5-fluoruracil (5-FU) and 5-fluoruridine (5-FUrd) was measured. The activity of these antibiotic/adjuvant combinations against MRSA growing planktonically and in biofilms was investigated and expanded to included OX. Finally the impact of reduced thymidine levels of accumulation of reactive oxygen species (ROS) and oxidative stress related membrane potential was investigated to gain further mechanistic insights. Our data reveal that the purine nucleoside adjuvant Gua also potentiates the activity of anti-folate antibiotics and pyrimidine analogues against MRSA. Furthermore enhanced adjuvant activity was measured when OX was included in these antibiotic/adjuvant combinations. These data further support the potential of Gua as an antibiotic adjuvant enhancing the activity of two separate classes of clinically used antibiotic against MRSA.

## Results

### The purine adjuvant guanosine downregulates thymidine levels revealing a new therapeutic target

To further understand the mechanistic basis for altered beta-lactam susceptibility in MRSA exposed to guanosine (Gua) or adenosine (Ade), intracellular metabolite levels were compared in cultures supplemented with either nucleoside alone or in combination with oxacillin (OX) (1 µg/ml) (Fig. S1 and S2). As expected, exposure to Gua or Ade was accompanied by significant intracellular accumulation of each nucleoside (Fig. S1), and pleiotropic effects on the metabolome (Fig. S2).

**Fig. S1.**
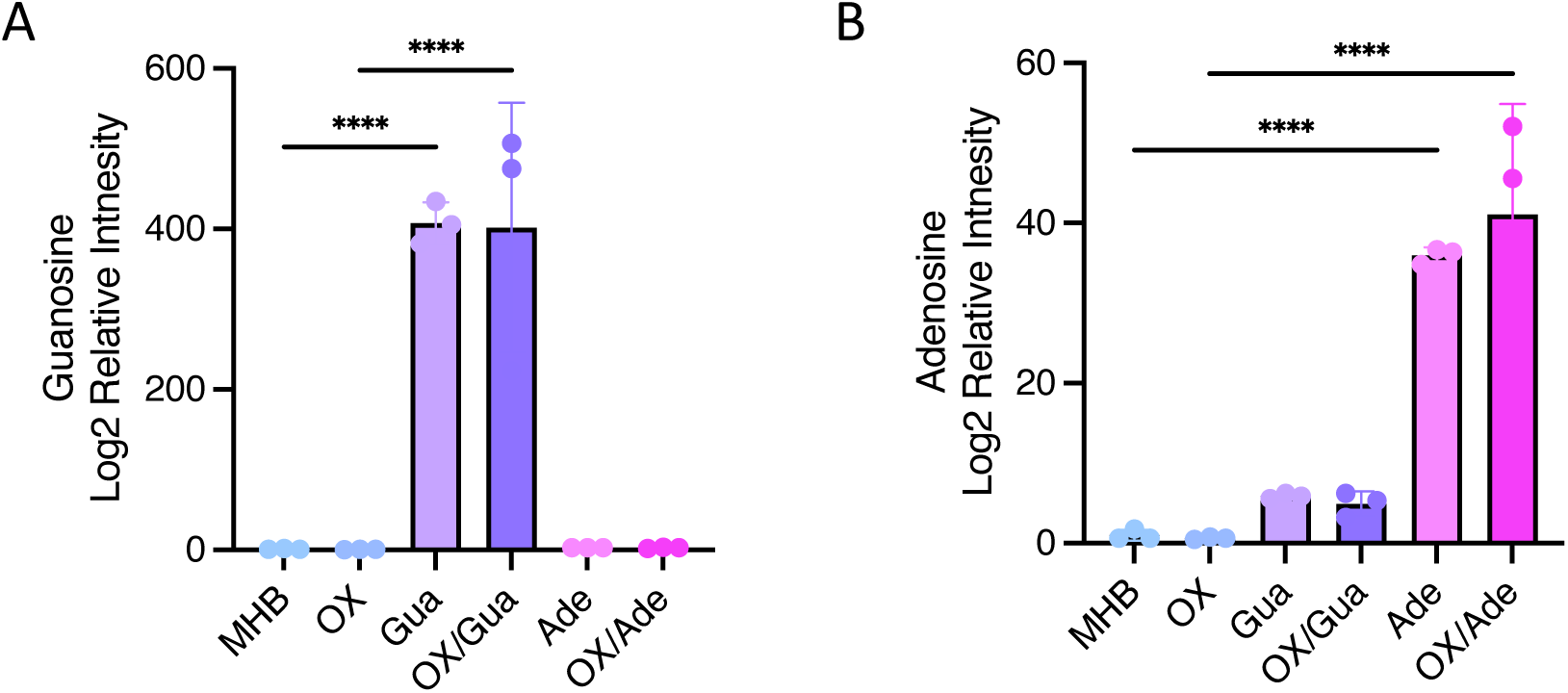
Impact of exogenous guanosine or adenosine alone and in combination with oxacillin on intracellular guanosine and adenosine in MRSA strain JE2. **A.** Intracellular guanosine (A) and adenosine (B) were quantified in cells grown in MHB, MHB oxacillin (OX, 1 µg/ml), MHB guanosine (Gua, 200 µg/ml), MHB OX/Gua, MHB adenosine (Ade, 200 µg/ml) and MHB OX/Ade. Data presented are the average of three biological replicates plotted using GraphPad Prism V9. Asterisks indicate statistically significant difference according to using a one-way ANOVA test. p-values **** p<0.0001 are indicated.

**Fig. S2.**
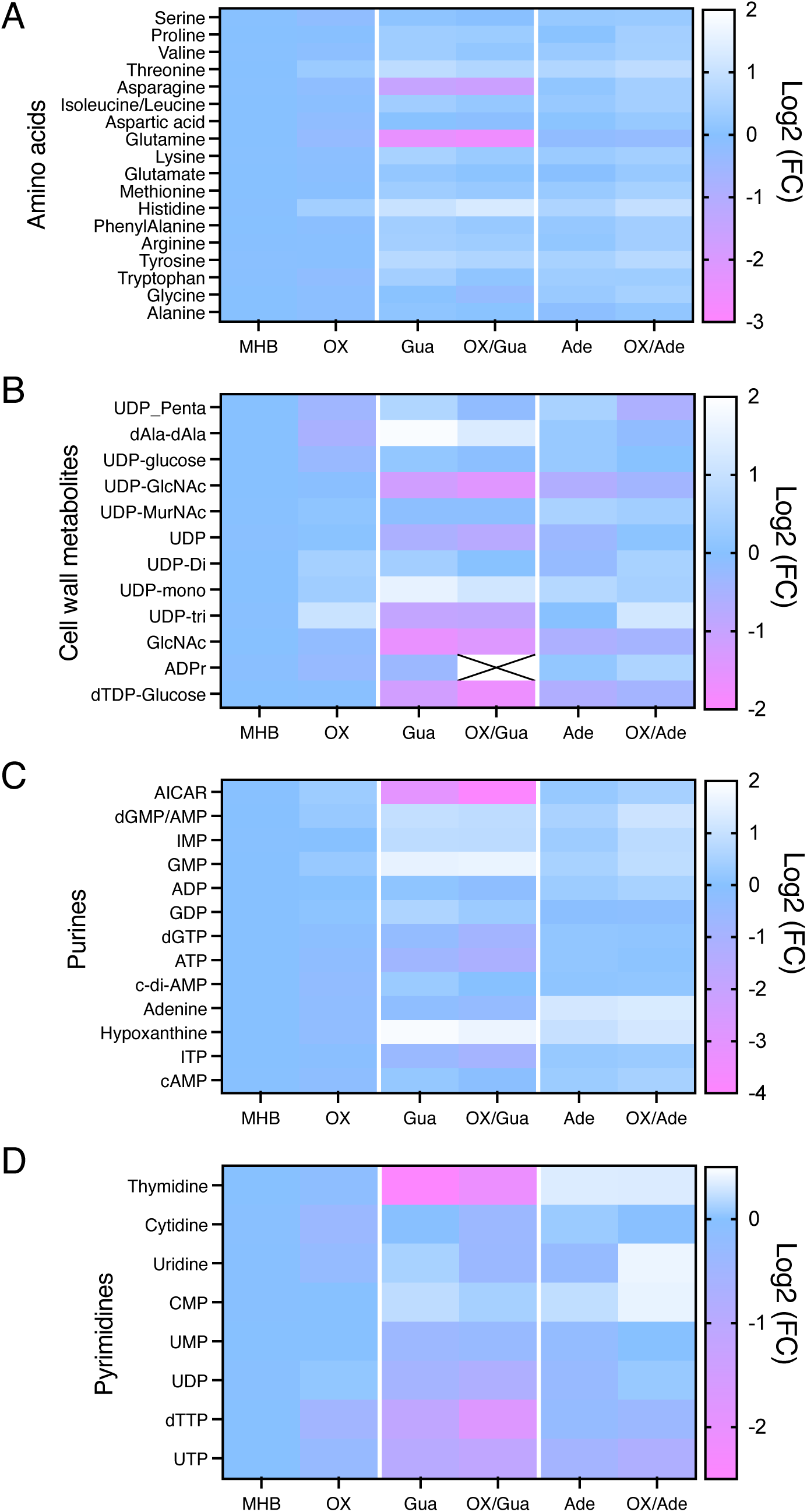
Exposure of MRSA to guanosine or adenosine has pleiotropic effects on the MRSA metabolome. Heatmap comparison of amino acids (A), cell wall metabolites (B), purines (C) and pyrimidines (D) in JE2 grown in MHB and MHB supplemented with guanosine (Gua, 200 µg/ml) or adenosine (Ade, 200 µg/ml) with or without oxacillin (OX, 1 µg/ml). The data presented are the average of three biological replicates analysed using GraphPad Prism V9. Individual metabolite levels that were significantly different were identified using a one-way ANOVA analysis.

Among the most notable changes were significant Gua-induced reductions in the levels of the purine biosynthetic intermediate AICAR, several cell wall precursors, the amino acids asparagine and glutamine and the pyrimidine nucleotide thymidine (Fig. S2). Asparagine serves as a precursor for purine biosynthesis (9), and its depletion may contribute to, or be a consequence of, disrupted purine homeostasis in MRSA treated with Gua. Glutamine is an important NH_4_^+^ donor for the amidation of iso-D-glutamate in the peptidoglycan stem pentapeptide (10). Furthermore glutamine is also an essential substrate for pyrimidine biosynthesis and its depletion in MRSA exposed to Gua may contribute to reduced levels of thymidine. The significant reduction in thymidine levels raised the possibility that Gua may also potentiate the activity of antibiotics, including anti-folates, that interfere with pyrimidine and/or thymidine biosynthesis (Fig. 1).

**Fig. 1.**
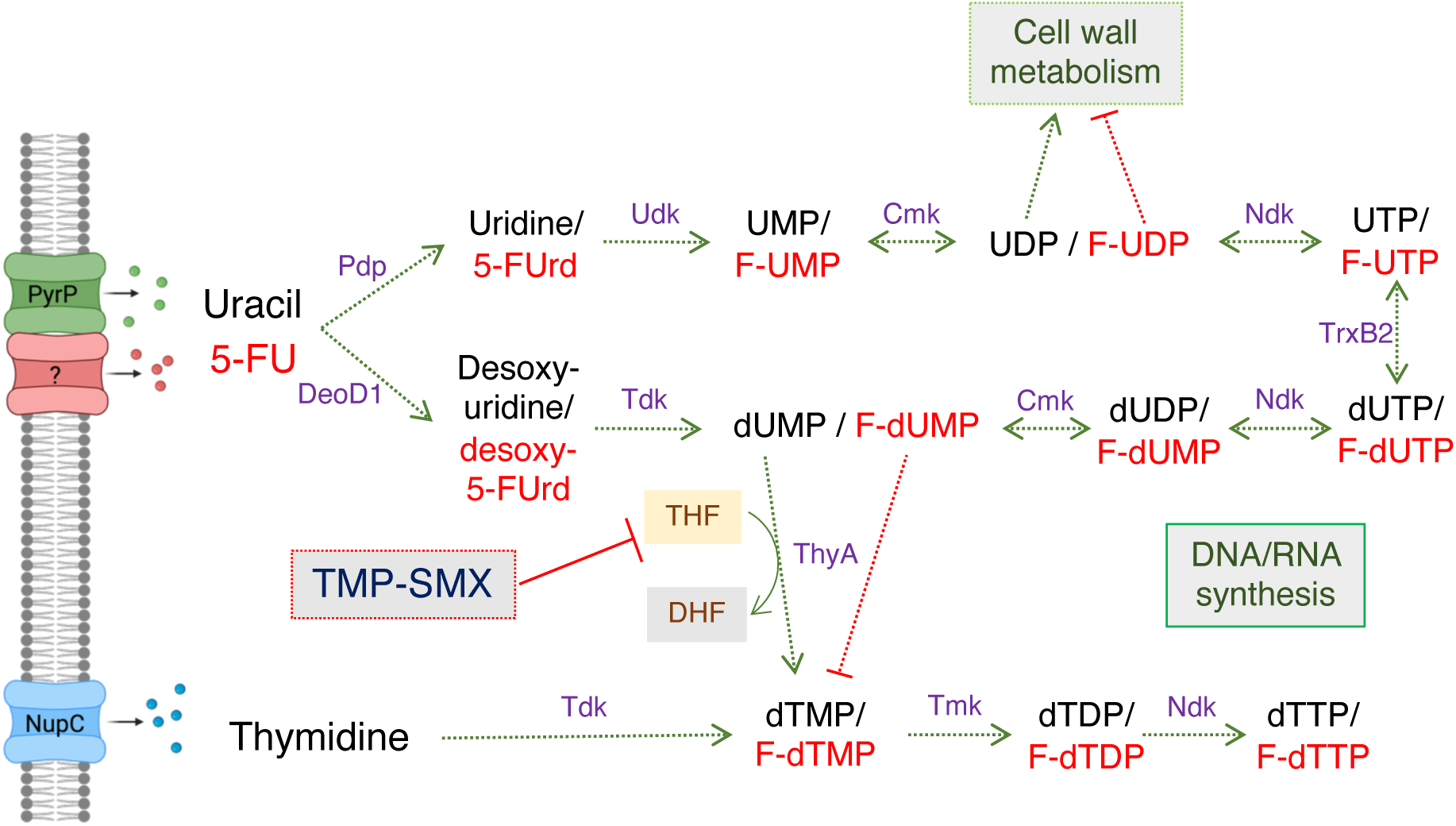
Overview of thymidine, uracil and 5-fluorouracil (5-FU) transport and metabolism including enzymes and intermediates implicated in susceptibility to 5-FU and trimethoprim-sulfamethoxazole (TMP-SMX). The permease PyrP (also known as UraA) has previously been implicated in transport of uracil (11, 12), but the transporter for 5-FU in *S. aureus* has not yet been identified. Uracil is converted to uridine and desoxy-uridine by Pdp and DeoD1, respectively, before being fluxed to UMP/dUMP, UDP/dUTP and UTP/dUTP. dUMP is converted to dTMP by thymidylate synthase (ThyA) (13) in a reaction requiring tetrahydrofolic acid (THF) as a co-factor, which is converted to dihydrofolic acid (DHF). UDP is an important intermediate in cell wall metabolism, which is adversely affected by the antimetabolite 5-F-UDP. 5-FU exposure leads to the build-up of F-UMP and F-TMP, which interferes with the normal incorporation of UMP and TMP into DNA and RNA, respectively. TMP-SMX inhibits two enzymatic reactions in the folate pathway thereby interfering with the synthesis of THF required for ThyA activity (7). Transport of exogenous thymidine by NupC (7) can bypass ThyA-dependent dTMP production. Figure created using biorender.com.

### Anti-MRSA activity of oxacillin combined with 5-fluorouracil or antifolates is significantly increased by guanosine

The impact of the Gua-induced reduction in thymidine levels on MRSA susceptibility to the anti-folate antibiotics trimethoprim-sulfamethoxazole (TMP-SMX) and the pyrimidine analogue drug 5-fluorouracil (5-FU) was investigated. 5-FU disrupts pyrimidine metabolism and has previously been reported to cause “thymidine-less death” in bacteria (14). Exogenous 5-FU is taken up and converted into fluorinated analogues of uridine/deoxyuridine; 5-fluorouridine/ 5-fluorodeoxyuridine (5-FUrd/deoxy 5-FUrd) (14–16) (Fig. 1). Checkerboard assays revealed significant synergy between Gua and 5-FU or 5-FUrd against JE2, but not with TMP, SMX or TMP-SMX (Table 1). Combinations of OX with 5-FU, 5-FUrd and TMP were also synergistic (Table 1). Strikingly, Gua (200 µg/ml) potentiated the activity of combinations of these antibiotics, as evidenced by lower ΣFIC values, including OX/SMX and OX/TMP-SMX that on their own were not synergistic (Table 1).

**Table 1.**
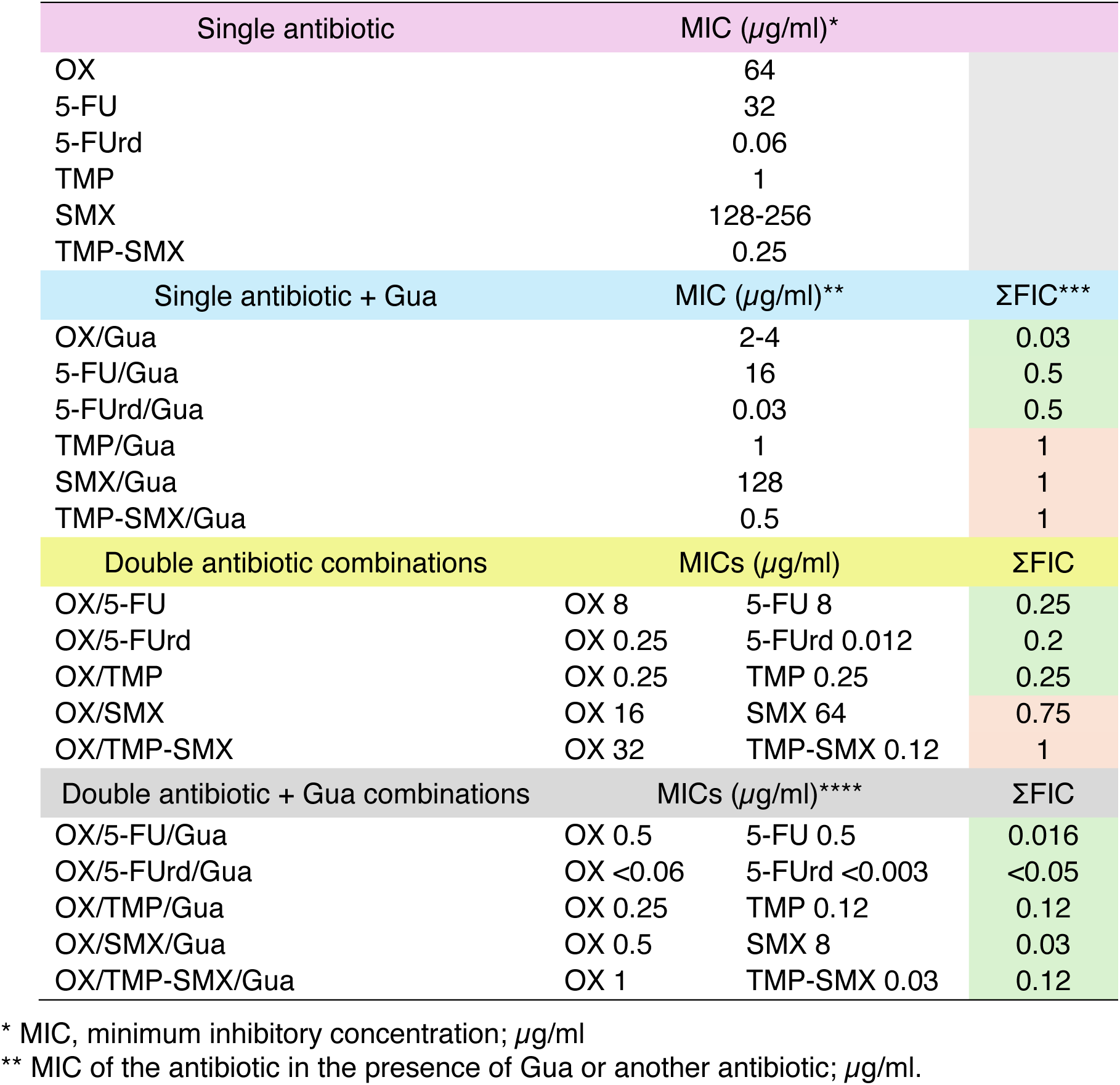
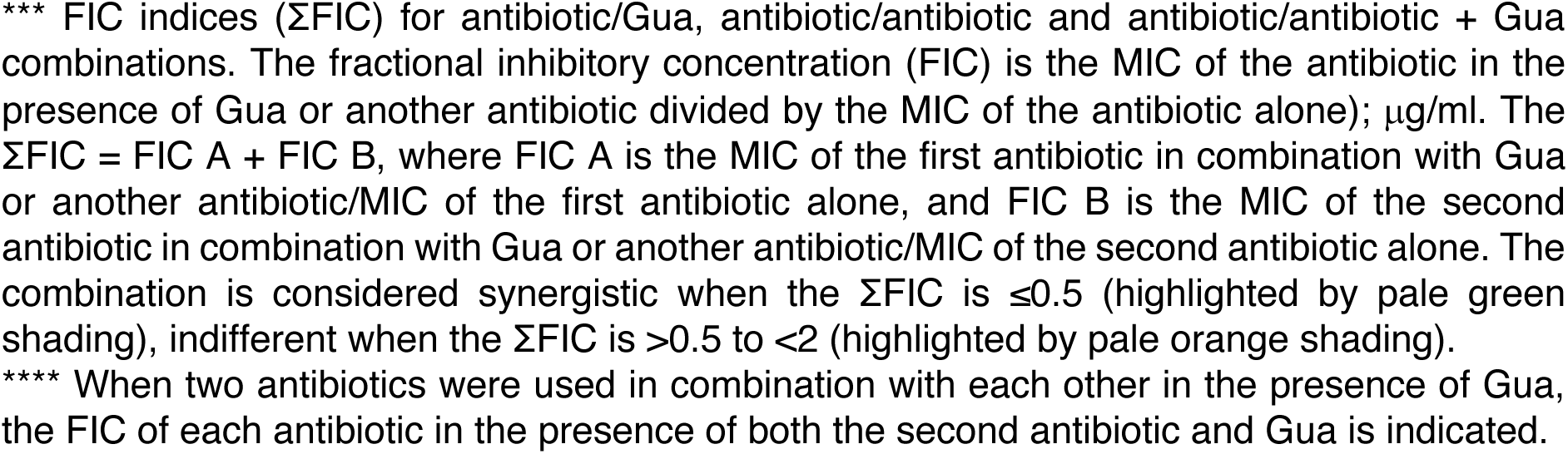
Antibacterial activity (minimum inhibitory concentrations, MIC) and drug synergy (fractional inhibitory concentration indices, ΣFIC) of oxacillin (OX), 5-fluorouracil (5-FU), 5-fluorouridine (5-FUrd), trimethoprim (TMP), sulfamethoxazole (SMX), TPM-SMX in combination with and without guanosine (Gua, 200 µg/ml), against MRSA strain JE2.

| Single antibiotic |  | MIC (μg/ml)* |  |  |
| --- | --- | --- | --- | --- |
| OX |  | 64 |  |  |
| 5-FU |  | 32 |  |  |
| 5-FUrd |  | 0.06 |  |  |
| TMP |  | 1 |  |  |
| SMX |  | 128-256 |  |  |
| TMP-SMX |  | 0.25 |  |  |
| Single antibiotic + Gua |  | MIC (μg/ml)** |  | ΣFIC*** |
| OX/Gua |  | 2-4 |  | 0.03 |
| 5-FU/Gua |  | 16 |  | 0.5 |
| 5-FUrd/Gua |  | 0.03 |  | 0.5 |
| TMP/Gua |  | 1 |  | 1 |
| SMX/Gua |  | 128 |  | 1 |
| TMP-SMX/Gua |  | 0.5 |  | 1 |
| Double antibiotic combinations |  | MICs (μg/ml) |  | ΣFIC |
| OX/5-FU | OX 8 | 5-FU 8 |  | 0.25 |
| OX/5-FUrd | OX 0.25 | 5-FUrd 0.012 |  | 0.2 |
| OX/TMP | OX 0.25 | TMP 0.25 |  | 0.25 |
| OX/SMX | OX 16 | SMX 64 |  | 0.75 |
| OX/TMP-SMX | OX 32 | TMP-SMX 0.12 |  | 1 |
| Double antibiotic + Gua combinations |  | MICs (μg/ml)**** |  | ΣFIC |
| OX/5-FU/Gua | OX 0.5 | 5-FU 0.5 |  | 0.016 |
| OX/5-FUrd/Gua | OX <0.06 | 5-FUrd <0.003 |  | <0.05 |
| OX/TMP/Gua | OX 0.25 | TMP 0.12 |  | 0.12 |
| OX/SMX/Gua | OX 0.5 | SMX 8 |  | 0.03 |
| OX/TMP-SMX/Gua | OX 1 | TMP-SMX 0.03 |  | 0.12 |

\* MIC, minimum inhibitory concentration; μg/ml
\*\* MIC of the antibiotic in the presence of Gua or another antibiotic; μg/ml.
\*\*\* FIC indices ( ΣFIC) for antibiotic/Gua, antibiotic/antibiotic and antibiotic/antibiotic + Gua combinations. The fractional inhibitory concentration (FIC) is the MIC of the antibiotic in the presence of Gua or another antibiotic divided by the MIC of the antibiotic alone); $\mu\text{g/ml}$ . The $\Sigma\text{FIC} = \text{FIC A} + \text{FIC B}$ , where FIC A is the MIC of the first antibiotic in combination with Gua or another antibiotic/MIC of the first antibiotic alone, and FIC B is the MIC of the second antibiotic in combination with Gua or another antibiotic/MIC of the second antibiotic alone. The combination is considered synergistic when the $\Sigma\text{FIC}$ is $\leq 0.5$ (highlighted by pale green shading), indifferent when the $\Sigma\text{FIC}$ is $>0.5$ to $<2$ (highlighted by pale orange shading).
\*\*\*\* When two antibiotics were used in combination with each other in the presence of Gua, the FIC of each antibiotic in the presence of both the second antibiotic and Gua is indicated.

Time-kill assays used to investigate potential bactericidal activity further revealed significant synergy (>2 log reduction in the number of JE2 colony forming units (CFUs)/ml) between Gua and 5-FU, 5-FUrd, SMX and TMP (0.5× MICs) (Fig. 2A-D), but not TMP-SMX (Fig. 2E). Notably bactericidal activity (> 3 log CFU/ml reduction) was achieved when low dose OX (4 µg/ml; 0.0625× MIC) was added to these antibiotic/Gua combinations (Fig. 2A-E). Further increasing the OX concentration to 32 µg/ml (0.5× MIC) achieved a near eradication of JE2 after 24 hours for all 5-FU, 5-FUrd, SMX, TMP and TMP-SMX combinations (Fig. 2F). Finally, the most significant bactericidal activity was measured when Gua was included with triple SMX/5-FUrd/OX, TMP/5-FU/OX, TMP/5-FUrd/OX and SMX/5-FU/OX combinations, which achieved eradication or near eradication of JE2 after 8-16 hr (Fig. 2G-J).

**Fig. 2.**
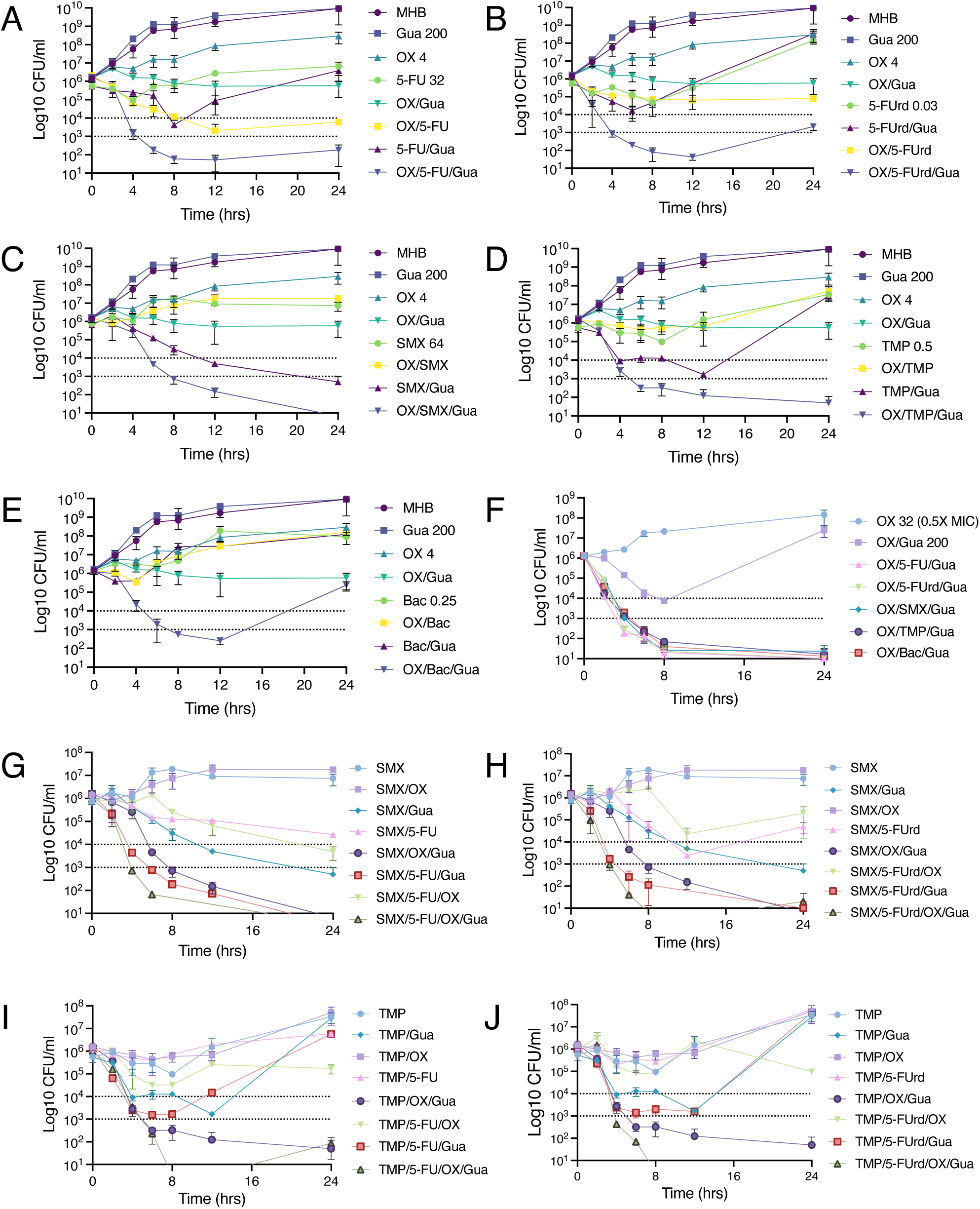
Guanosine potentiates MRSA killing by oxacillin, anti-folate antibiotics, and pyrimidine analogues. Time-kill assays were performed using 0.5× MIC concentrations of 5-fluorouracil (5-FU; 32 µg/ml), 5-fluorouridine (5-FUrd; 0.03 µg/ml), sulfamethoxazole (SMX; 64 µg/ml), trimethoprim (TMP; 0.5 µg/ml), or Bactrim (SMX-TMP; 0.25 µg/ml), alone or in combination with guanosine (Gua; 200 µg/ml) and/or oxacillin (OX; 4 µg/ml [0.0625× MIC] or 32 µg/ml [0.5× MIC]). Panels show killing kinetics for: **A.** 5-FU ± OX/Gua; **B.** 5-FUrd ± OX/Gua; **C.** SMX ± OX/Gua; **D.** TMP ± OX/Gua; **E.** Bactrim ± OX/Gua; **F.** 0.5× OX ± Gua or Gua/antimetabolite combinations; **G.** SMX ± OX, 5-FU and Gua, **H.** SMX ± OX, 5-FUrd and Gua; **I.** TMP ± OX, 5-FU and Gua and **J.** TMP ± OX, 5-FUrd and Gua. Exponential phase cultures were inoculated into MHB 2% NaCl at a starting cell density of approximately 1×10^6^ CFU/ml, with or without antibiotics and Gua as indicated, and CFUs enumerated after 0, 2, 4, 6, 8, 12, and 24 h. The data presented are the average of three independent experiments plotted using GraphPad Prism V9 and standard deviations are shown.

Using Nebraska Transposon Mutant Library (NTML) mutants implicated in nucleotide transport and metabolism revealed that the reversal of 5-FU- and 5-FUrd-induced JE2 growth inhibition by exogenous thymidine or uridine, respectively (Fig. S3), was *nupC*-dependent. NupC was previously reported as the primary thymidine transporter in *S. aureus* (7).

Extending these analyses to other MRSA strains, disk diffusion assays revealed that Gua also potentiated the activity of 5-FU and 5-FUrd, but not SMX or TMP, against MW2, COL and BH1CC (Table S1). However double disk diffusion assays with SMX and TMP also revealed that synergy between these two anti-folates against MW2, COL and BH1CC was enhanced by Gua (Table S1). Furthermore the anti-MRSA activity of combinations of these antimicrobial drugs was further enhanced by Gua for all strains (the only exception was SMX-5-FU/Gua against MW2) (Table S1).

Taken together, these findings support an important role for the down-regulation of thymidine in the antibiotic adjuvant activity of Gua. Gua potentiates the anti-MRSA activity of anti-folate and pyrimidine analogue drugs, as well as β-lactams. Combinations of these antibiotics with Gua can eradicate MRSA, albeit in a strain-dependent manner.

**Fig. S3.**
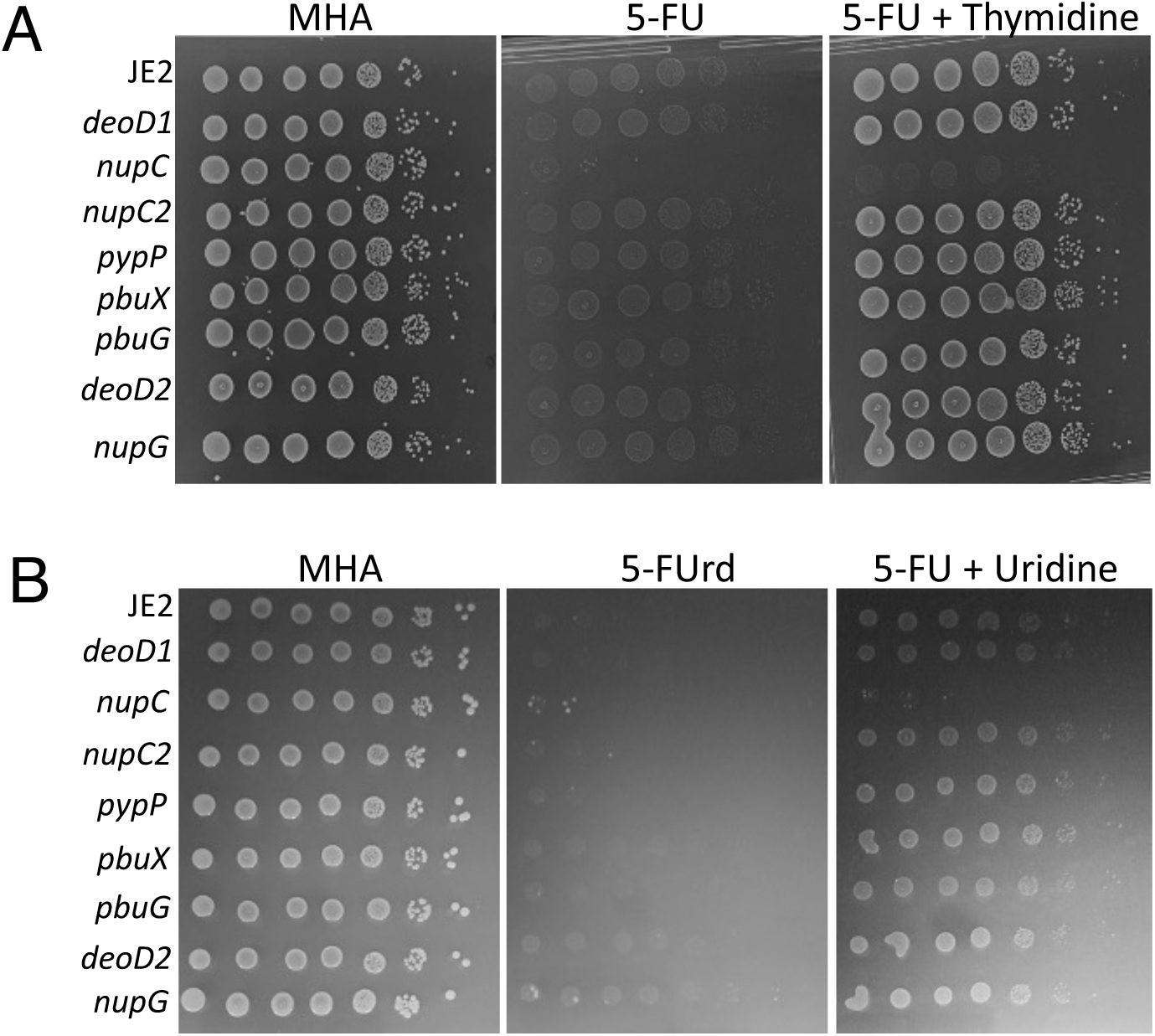
Thymidine or uridine rescue MRSA growth inhibition by 5-fluorouracil (A) and 5-fluoruridine (B), respectively, in a *nupC*-dependent manner. Serial dilution spot plate assays using NTML mutants implicated in nucleotide transport and metabolism on Mueller-Hinton agar (MHA), MHA supplemented with 5-fluorouracil (5-FU; 8 µg/ml) or 5-fluorouridine (5-FUrd; 1 µg/ml), and MHA containing 5-FU or 5-FUrd plus thymidine or uridine (50 µg/ml). Plates were incubated for 24 h. These experiments were repeated 3 times, and representative plates are shown.

**Table S1.** Antibacterial activity (zone diameters, mm) and synergy of cloxacillin (Clox), sulfamethoxazole (SMX), trimethoprim (TMP), 5-fluorouracil (5-FU), 5-fluorouridine (5-FUrd), and Bactrim (Bac), alone and in combination with guanosine (Gua, 200 µg/ml), against MRSA strains JE2, MW2, COL, and BH1CC.

| Drug +/- Gua | Strain | JE2 | MW2 | COL | BH1CC |
| --- | --- | --- | --- | --- | --- |
| Clox |  | 11 | 6 | 6 | 6 |
| Clox/Gua |  | 29 | 12 | 6 | 9 |
| SMX |  | 6 | 6 | 6 | 6 |
| SMX/Gua |  | 15 | 6 | 6 | 14 |
| TMP |  | 19 | 18 | 17 | 19 |
| TMP/Gua |  | 20 | 20 | 21 | 21 |
| 5-FU |  | 14 | 6 | 11 | 19 |
| 5-FU/Gua |  | 17 | 12 | 25 | 19 |
| 5-FUrd |  | 15 | 11 | 16 | 16 |
| 5-FUrd/Gua |  | 16 | 17 | 24 | 20 |
| Double drug combinations +/- Gua<sup>1</sup> |  |  |  |  |  |
| SMX-TMP |  | + | + | + | + |
| SMX-TMP/Gua |  | ++ | (+) | ++ | (+) |
| Clox-5-FU |  | + | - | + | - |
| Clox-5-FU/Gua |  | ++ | ++ | ++ | ++ |
| Clox-5-FUrd |  | + | + | - | - |
| Clox-5-FUrd/Gua |  | ++ | ++ | ++ | ++ |
| SMX-5-FU |  | + | - | - | - |
| SMX-FU/Gua |  | ++ | - | ++ | ++ |
| SMX-5-FUrd |  | + | - | - | - |
| SMX-5-FUrd/Gua |  | ++ | ++ | ++ | ++ |
| TMP-5-FU |  | + | - | + | + |
| TMP-5-FU/Gua |  | (+) | ++ | ++ | ++ |
| TMP-5-FUrd |  | + | + | + | + |
| TMP-5-FU/Gua |  | (+) | ++ | ++ | ++ |

<sup>1</sup> Synergy was assessed by placing two antibiotic-impregnated disks 10 mm apart on MHA plates and defined as an increased zone of inhibition in the region between the disks relative to single-drug controls. “-“ no detectable synergy; “+” synergy in the absence of Gua; “(+)” no increase in synergy in the presence of Gua; “++” enhanced synergy between the two drugs in the presence of Gua.

### Eradication of MRSA biofilms by antibiotic/guanosine combinations

Antibiotics were tested at concentrations approximating their reported maximum serum levels (C_max_): oxacillin (OX, 200 µg/ml) (17–19), sulfamethoxazole (SMX, 78 µg/ml) and trimethoprim (TMP, 4 µg/ml) (20), 5-fluorouracil (5-FU, 68 µg/ml) (18), 5-fluorouridine (5-FUrd, 200 µg/ml) (21), and vancomycin (VAN, 50 µg/ml) (22), which is used to treat MRSA infections was included as a positive control. Each antibiotic was evaluated alone and in combination with guanosine (Gua, 200 µg/ml), which is not reported to be toxic (23).

Exposure of pre-formed JE2 biofilms to OX alone produced a ∼2-fold reduction in CFU/ml, comparable to VAN. The OX/Gua combination further improved activity, achieving a ∼3-fold reduction (Fig. 3). TMP and SMX, individually or combined with Gua, showed minimal anti-biofilm activity. In contrast, 5-FU and 5-FUrd were more effective, particularly when paired with Gua (Fig. 3). Strikingly, combinations of OX with TMP, SMX, 5-FU, or 5-FUrd yielded 2–4 log reductions in CFU/ml (Fig. 3). Although adding Gua did not enhance the activity of OX/5-FU or OX/5-FUrd, both OX/5-FUrd and OX/5-FUrd/Gua achieved 4–5 log reductions, significantly greater than the 2-log reduction observed with VAN (Fig. 3) and surpassing the 1-3 log reductions typically reported for single antimicrobial agents (24). These 4-5 log reductions are in line with accepted thresholds for disinfection and are comparable to the most effective anti-biofilm antimicrobial combinations reported to date (24–26).

**Fig. 3.**
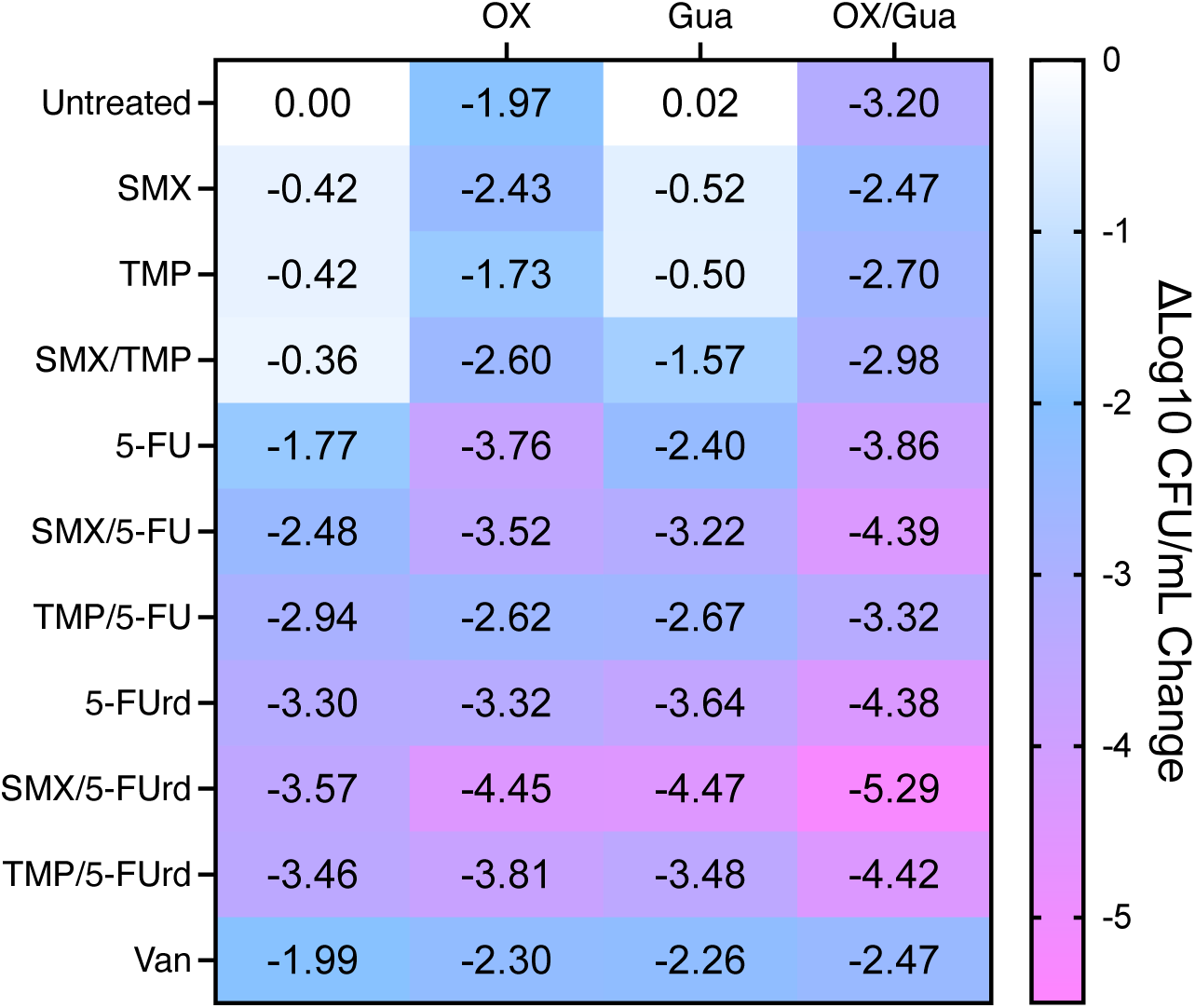
Oxacillin/guanosine combinations with anti-folates and pyrimidine analogues enhance JE2 biofilm eradication. Pre-formed 24h JE2 biofilms were treated for an additional 24h with antibiotics or the adjuvant guanosine singly and in combinations at concentrations approximating reported Cmax values. Surviving cells were serially diluted, enumerated on MHA, and the results presented as Δlog₁₀ CFU/ml relative to untreated controls. The antibiotic/adjuvant concentrations used were: OX, 200 μg/ml; SMX, 78 μg/ml; TMP, 4 μg/ml; 5-FU, 68 μg/ml; 5-FUrd, 200 μg/ml; VAN, 50 μg/ml and Gua, 200 μg/ml. The data, which represent the mean of 3 independent experiments, were plotted using GraphPad Prism v9.

### Guanosine exacerbates cell envelope stress induced by oxacillin and 5-fluorouracil

Confocal microscopy imaging of JE2 exposed to OX, 5-FU and Gua, alone and in combinations was used to probe the mechanism(s) of growth inhibition and cell killing. Gua alone did not alter membrane staining with FM4-64 (27) or peptidoglycan incorporation visualized with HADA (28, 29). As expected, HADA fluorescence localized predominantly at the division septum in untreated and Gua-treated cells (Fig. 4A, E). Consistent with previous reports (1, 30–33), OX treatment resulted in visibly enlarged cells with characteristic membrane bulges at sites perpendicular to the septum (Fig. 4B). Subinhibitory 5-fluorouracil (5-FU) resulted in more diffuse HADA labelling throughout the cell wall, indicating altered peptidoglycan synthesis compared to untreated or Gua-treated controls (Fig. 4C).

**Fig. 4.**
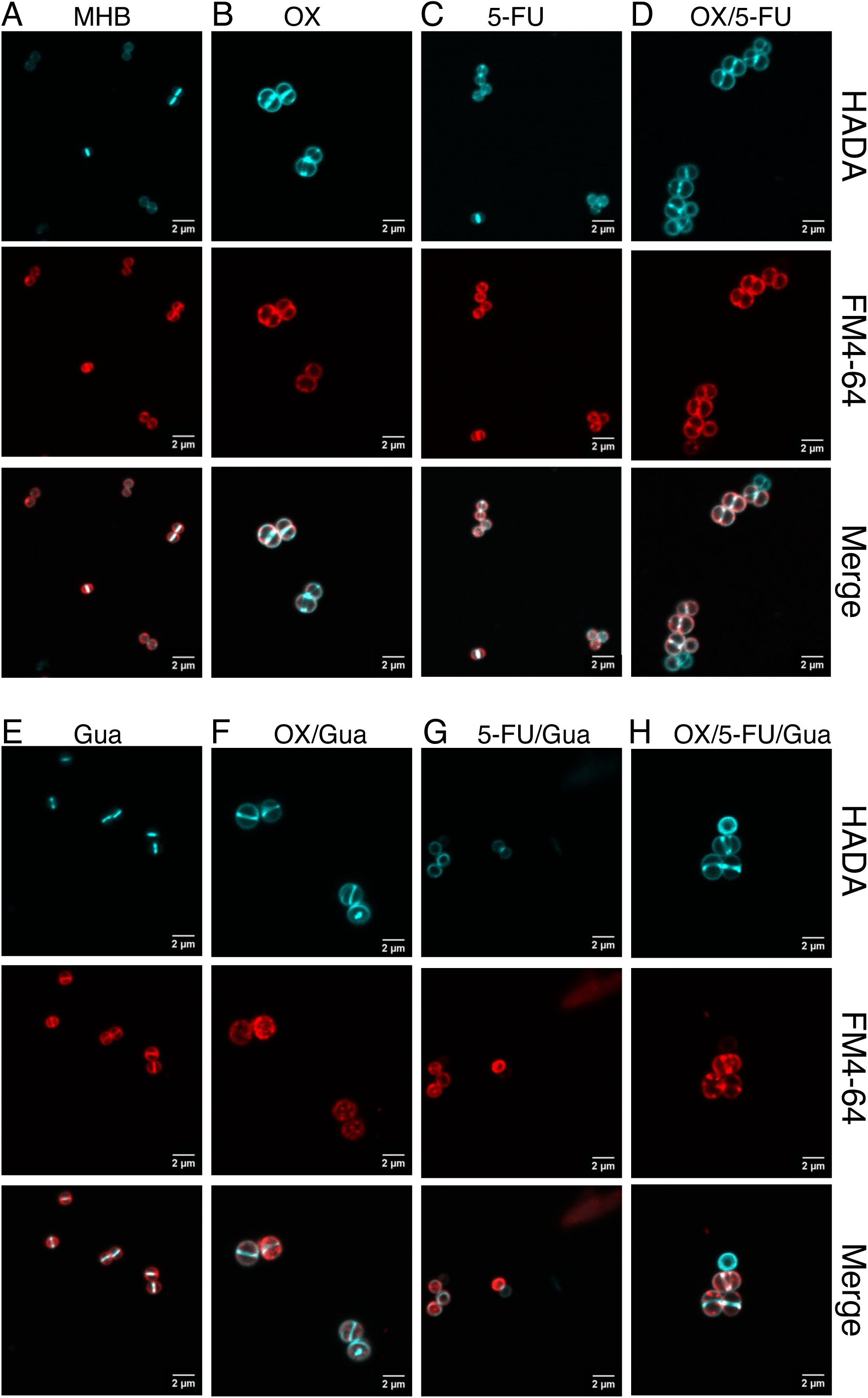
Guanosine potentiates oxacillin/5-fluorouracil-induced perturbation of the MRSA cell envelope. Representative confocal microscopy images of JE2 cells grown in: **A.** MHB; **B.** MHB + oxacillin (OX, 1 µg/ml); **C.** MHB + guanosine (Gua, 200 µg/ml); **D.** MHB + OX/Gua; **E.** MHB + 5-fluorouracil (5-FU, 1 µg/ml); **F.** MHB + 5-FU/Gua; **G.** MHB + OX/5-FU; **H.** MHB + OX/5-FU/Gua. Cells were stained with HADA to label newly synthesized peptidoglycan and FM4-64 to visualize *S. aureus* membranes. White arrows highlight cells that incorporated HADA but did not take up FM4-64. Yellow triangles mark cells with membrane dye aggregates, and white triangles indicate cells exhibiting HADA-labelled peptidoglycan clumps. Images were acquired at 1000× magnification using an Olympus LS FLUOVIEW FV3000 confocal laser scanning microscope and processed with ImageJ (Fiji v1.0).

Cells treated with the OX/5-FU combination resembled OX-treated cells in both HADA and FM4-64 staining patterns (Fig. 4D). However, closer examination revealed that some OX/5-FU-treated cells incorporated HADA while failing to incorporate FM4-64, suggesting altered membrane dye uptake despite normal peptidoglycan staining.

Cell staining was more unpredictable when Gua was combined with OX, 5-FU, or both (Fig. 4F-H). OX/Gua-treated cells exhibited prominent HADA- and FM4-64-strained intracellular clumps that were absent from cells treated with OX alone (Fig. 4F), suggestive of perturbed peptidoglycan synthesis and abnormal membrane organization. In 5-FU/Gua-treated cells, HADA incorporation remained detectable, but FM4-64 labelling was frequently absent or restricted to irregular membrane clusters, resembling patterns previously associated with weakened cell wall integrity (34) (Fig. 4G).

The triple combination (OX/5-FU/Gua) produced the most severe defects, including cells lacking FM4-64 staining altogether, extensive membrane clumping, and multiple HADA-labelled division sites (Fig. 4H). These features indicate simultaneous disruption of membrane architecture and septal peptidoglycan synthesis.

Taken together, these observations show that while OX and 5-FU individually perturb distinct aspects of the cell envelope, the addition of Gua intensifies these defects, generating cells with inconsistent membrane dye uptake, aberrant peptidoglycan accumulation, and multiple or stalled division sites. The combined OX/5-FU/Gua treatment therefore exerts pleiotropic and synergistic effects on both membrane integrity and cell wall synthesis, consistent with extensive cell envelope stress.

### Guanosine exacerbates ROS accumulation and disrupts membrane potential in combination with oxacillin and 5-fluorouracil

Reactive oxygen species (ROS) contribute to the mechanism of action for compounds targeting thymidine biosynthesis including SMX-TMP (35, 36). ROS accumulation may in turn impact membrane integrity as described earlier (Fig. 5). ROS levels were quantified using H_2_DCFDA, a fluorogenic probe activated by intracellular oxidation (37). Cells were grown to exponential phase, treated with OX, Gua, 5-FU, or their combinations, and analysed by flow cytometry alongside an H2O2 control. OX treatment increased ROS levels (Fig. 5A), consistent with β-lactam-induced ROS accumulation (38). ROS levels in cells treated with OX/5-FU were similar to OX alone, whereas 5-FU alone had little effect. Gua-treated cells also showed elevated ROS. Strikingly, OX/Gua and OX/Gua/5-FU combinations produced ROS levels comparable to the strong inducer H2O2 (39), indicating severe oxidative stress under these conditions. In killing assays, the ROS scavenger glutathione (10 mM) (40) reversed the bactericidal activity of antibiotic (OX, SMX, TMP, Bac, 5-FU and 5-FUrd) / Gua combinations (Fig. 5B-G), supporting the importance of ROS in the adjuvant activity of Gua. Glutathione alone did not affect MRSA killing by any of these antibiotics (Fig. 5B-G).

**Fig. 5.**
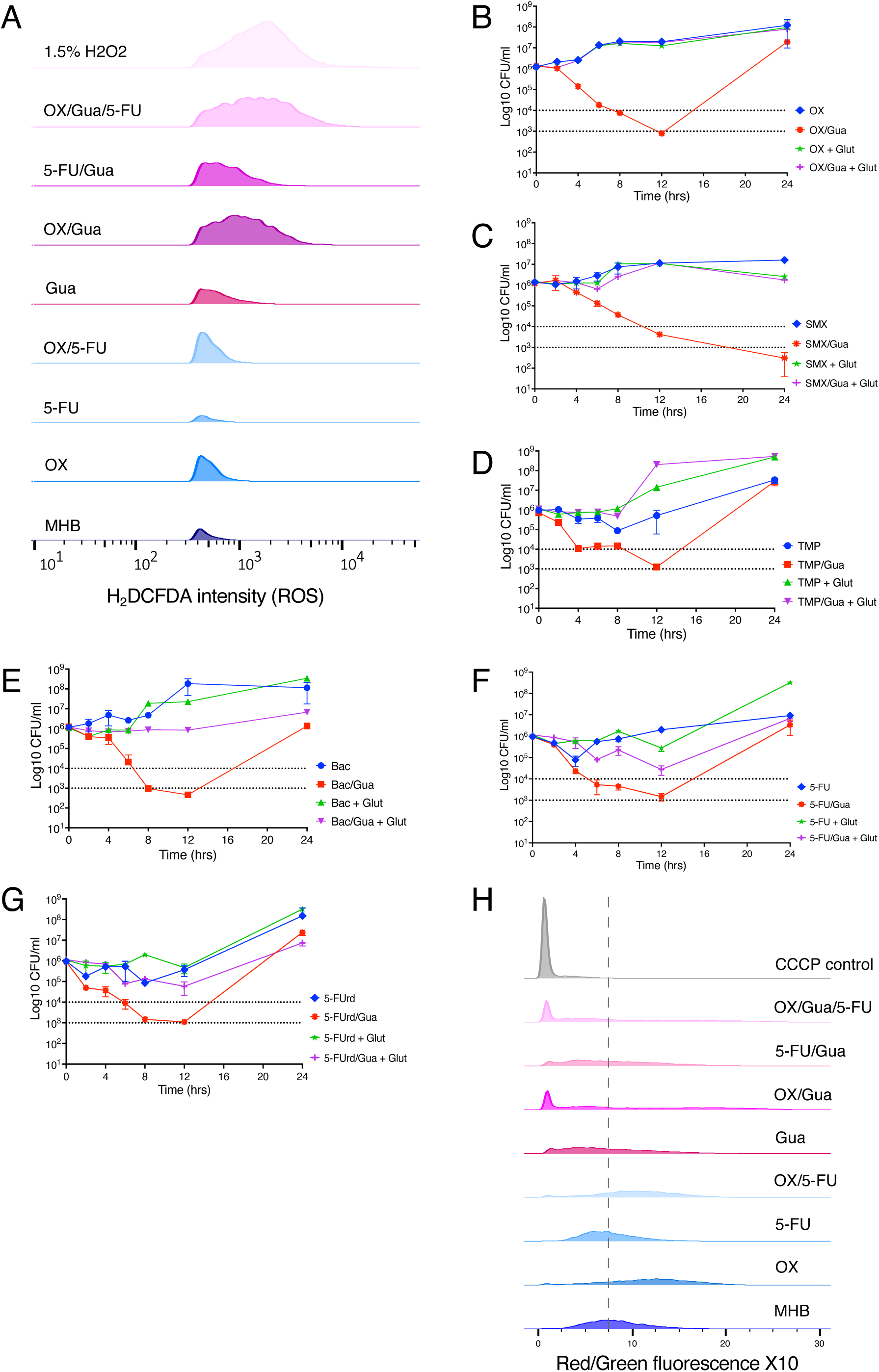
Guanosine enhances antibiotic-induced oxidative stress and membrane depolarization in MRSA. **A.** Intracellular ROS levels measured using the fluorescent probe carboxy-H2DCFDA. JE2 cells were grown to exponential phase and exposed to Gua, OX, 5-FU, or their combinations, then stained with carboxy-H2DCFDA (10 µM) for 10 min. A 1.5% H2O2 treatment served as a positive control and produced robust fluorescence. Histograms show relative fluorescence versus cell counts. Data were processed using FlowJo v10.1 and one biological replicate from three independent experiments is shown. **B.-G.** Time-kill assays comparing the impact of guanosine and antioxidant supplementation on antibiotic-mediated killing. Exponential phase cultures were treated with oxacillin (OX, 32 µg/ml), sulfamethoxazole (SMX, 64 µg/ml), trimethoprim (TMP, 0.5 µg/ml), Bactrim (Bac, 0.25 µg/ml), 5-fluorouracil (5-FU, 32 µg/ml), 5-fluorouridine (5-FUrd, 0.06 µg/ml) alone or in combination with guanosine (Gua; 200 µg/ml) and/or glutathione (Glut, 10mM). Panels show killing kinetics for: **B.** OX ± Gua alone or Gua + Glut. **C.** SMX ± Gua alone or Gua + Glut. **D.** TMP ± Gua alone or Gua + Glut. **E.** Bac ± Gua alone or Gua + Glut. **F.** 5-FU ± Gua alone or Gua + Glut. **G.** 5-FUrd ± Gua alone or Gua + Glut. Exponential phase cultures were inoculated into MHB 2% NaCl at a starting cell density of approximately 1×10^6^ CFU/ml, with or without antibiotics and Gua as indicated, and CFUs enumerated after 0, 2, 4, 6, 8, 12, and 24 h. The data presented are the average of three independent experiments plotted using GraphPad Prism V9 and standard deviations are shown. **H.** Membrane potential measurements using DiOC2(3) and flow cytometry. Exponential-phase JE2 cells were treated with Gua, OX, 5-FU, or their combinations, stained with DiOC2(3), and analysed by FACS. Carbonyl cyanide m-chlorophenylhydrazone (CCCP) (15 µM final concentration) was used as a control to collapse the membrane potential. For each sample, 10,000 gated events were collected. Red/green fluorescence ratios ×10 were plotted as histograms. Data were processed using FlowJo v10.1 and one biological replicate from four independent experiments is shown.

Membrane potential was assessed using the fluorescent dye DiOC2(3) (41). Increased red:green fluorescence ratios indicate elevated membrane potential, whereas decreased ratios reflect dissipation of the proton motive force (PMF), a hallmark of cell death (42, 43). OX increased the red:green ratio, while 5-FU had minimal effect and OX/5-FU resembled OX-treated cells (Fig. 5H). In contrast, Gua-treated cells displayed a reduced red:green ratio, indicating membrane depolarization. Importantly, Gua/OX and Gua/OX/5-FU treatments were accompanied by a pronounced shift toward low red:green ratios, comparable to carbonyl cyanide m-chlorophenyl hydrazone (CCCP)-treated cells, indicative of a substantial PMF collapse that may contribute to reduced viability (Fig. 5H). Together, these data indicate that Gua-induced ROS and membrane depolarisation, which are amplified in combination with OX (and to a lesser extent 5-FU) are likely contribute to extensive cell envelope defects, growth inhibition and cell killing.

## Discussion

Building on our recent report that the purine nucleosides guanosine and xanthosine increase MRSA susceptibility to beta-lactam antibiotics (1), we report here that exposure to guanosine significantly reduces thymidine levels and increases susceptibility to the anti-folate antibiotics SMX and TMP and the pyrimidine analogues. 5-FU and 5-FUrd. SMX and TMP are cost-effective antibiotics often used in combination for the treatment of MRSA infections. These drugs target sequential enzymes in folate synthesis that feed into thymidine production (44–46). but their effectiveness is undermined by resistance associated with mutations in *dfrB* and *thyA* (8, 47–49). The efficacy of SMX or TMP ultimately depends on thymidine levels (46) and growth inhibition of JE2 on blood agar plates supplemented with TMP was previously shown to be thymidine dependent (13). Our data reveal potent anti-MRSA activity of either SMX or TMP in combination with both Gua and OX. This novel drug combination may also limit the emergence of SMX-TMP resistant MRSA strains and be effective against MRSA strains resistant to non-β-lactam antimicrobial drugs.

Reductions in the levels of the purine intermediate AICAR, glutamine and thymidine were among the most striking observations from metabolomic analysis of MRSA exposed to guanosine. Interestingly, levels of the pyrimidine nucleotide cytidine were not significantly reduced by Gua. The major impact of guanosine on nucleotide homeostasis makes it challenging to speculate on why glutamine and thymidine levels in particular are significantly downregulated. Because thymidine, but not cytidine, biosynthesis depends on tetrahydrofolate (THF), the reduced levels of thymidine in Gua-treated cells is indicative of impaired folate metabolism. GTP is the substrate for the first committed step of folate biosynthesis, the FolE/FolE2-catalyzed conversion of GTP to dihydroneopterin triphosphate (50, 51). Elevated GMP levels (Fig. S2) in Gua-treated cells may also interfere with normal levels of purine biosynthesis feedback inhibition, thereby impacting purine/GTP-dependent THF availability and thymidine biosynthesis. Reduced levels of several peptidoglycan precursors in Gua-treated cells may also be a consequence of reduced glutamine availability, and contribute to enhanced β-lactam susceptibility. MRSA cells treated with guanosine and oxacillin also displayed altered membrane labelling using FM4-64, an observation that may also be consistent with downregulation of peptidoglycan synthesis enzymes and beta-lactam resistance (52–54).

The data presented here indicate that reduced thymidine in OX/Gua-treated MRSA exposes an enhanced vulnerability to pyrimidine antimetabolites (5-FU and 5-FUrd) that target thymidine synthesis (55). Guanosine-mediated depletion of thymidine also potentiates the activity of SMX, TMP, and the SMX-TMP combination when used with oxacillin revealing an alternative treatment strategy for infections typically managed with SMX-TMP alone, particularly where therapeutic failures or suboptimal responses arise (8, 56). For example SMX-TMP has relatively weak activity against MRSA biofilms (57) whereas the guanosine/oxacillin/5-FU or 5-FUrd combinations described in this study also have significant anti-biofilm activity raising the possibility that they could be used to improve the management of device-related and other chronic infections. 5-FU has previously been reported to interfere with LuxS/autoinducer-2-regulated biofilm in *S. aureus* (58). The antibacterial properties of fluorinated pyrimidine antagonists, which are used in the treatment of cancer over longer time periods, indicates their potential clinical usefulness as antimicrobial drugs (59, 60). Indeed, 5-FU and 5-FUrd have been used successfully to treat staphylococcal infections in mice (61, 62). The absence of cross resistance between 5-FU/5-FUrd and SMX/TMP further paves the way to their deployment in the treatment of infections caused by TMP-SMX resistant *S. aureus* (63).

The consequences of the guanosine-induced reduction in thymidine levels were evident in both increased ROS generation and dissipated membrane potential. ROS is a known contributor to the mechanism of action of compounds such as SMX-TMP that reduce thymidine levels and is accompanied by DNA damage and cell death (35, 36). Levels of ROS in MRSA exposed to guanosine or guanosine/oxacillin/5-FU combinations were comparable to those induced by 1.5% H_2_O_2_. Supporting a role for oxidative stress in bactericidal activity, the ROS scavenger glutathione attenuated MRSA killing by antibiotic/guanosine combinations. Increased ROS levels were also accompanied by dissipation of membrane potential in cells treated with oxacillin and guanosine, suggesting that oxidative damage compromises proton motive force and membrane energetics. ROS-driven collapse of membrane potential has previously been linked to bacterial cell death (64), and may also contribute to enhanced MRSA killing.

Together, these data reveal a novel therapeutic approach using the adjuvant guanosine to re-sensitize MRSA to β-lactams and clinically used anti-folate or pyrimidine-based drugs, supporting the development of combination therapies that improve the treatment options for MRSA infections without requiring new classes of antimicrobial drug.

## Materials and Methods

### Bacterial strains and growth conditions

Bacterial strains are listed in Table S2. All strains were grown in Mueller-Hinton Broth (MHB) or Mueller-Hinton Agar (MHA), supplemented with guanosine (Gua), adenosine (Ade), 5-fluorouracil (5-FU) where indicated.

**Table S2.** Bacterial strains and plasmids used in this study

| Strains | Relevant Details |
| --- | --- |
| JE2 | USA300 cured of p01 & p03. Parent of Nebraska Transposon Mutant Library (NTML). |
| MW2 | MRSA SCCmec type IV; CC1 (65) |
| COL | MRSA reference strain; SCCmec type I; CC8 (66) |
| BH1CC | MRSA clinical isolate; SCCmec type II; CC8 (67) |
| NE1419 *nupG* | JE2 *nupG* (SAUSA300_0611). Erm<sup>r</sup>. (68) |
| NE283 *pbuG/stgP* | JE2 *pbuG/stgP* (SAUSA300_2207). Erm<sup>r</sup>. (68) |
| NE280 *pbuX* | JE2 *pbuX* (SAUSA300_0387). Erm<sup>r</sup>. (68) |
| NE650 *deoD2* | JE2 *deoD2* (SAUSA300_2091). Erm<sup>r</sup>. (68) |
| NE477 *deoD1* | JE2 *deoD1* (SAUSA300_0138). Erm<sup>r</sup>. (68) |
| NE544 *nupC1* | JE2 *nupC1* (SAUSA300_0298). Erm<sup>r</sup>. (68) |
| NE622 *nupC2* | JE2 *nupC2* (SAUSA300_0313). Erm<sup>r</sup>. (68) |
| NE1048 *pyrP* | JE2 *pyrP* (SAUSA300_1092). Erm<sup>r</sup>. (68) |

### Comparative metabolomic analysis

Intracellular metabolite analysis was performed as described previously (69, 70). 250 ml flasks containing 25 ml MHB (with or without 1 μg/ml OX, 200 µg/ml Gua or 200 µg/ml Ade) were inoculated from 5 ml overnight cultures at a starting OD_600_=0.06 and grown for 4–5 h. Culture volumes corresponding to OD_600_=10 were harvested and rapidly filtered through a 0.45 µm membrane (Millipore). The cells on the membrane were washed twice with 5 ml cold saline and immediately quenched in ice-cold 60% ethanol containing 2 µM Br-ATP and 2 µM Ribitol as internal controls. The cells were mechanically disrupted using a bead homogenizer set to oscillate for 3 × 30 s cycles at 6800 rpm with a 10 s pause between each cycle. Cell debris was separated by centrifugation at 12,000 rpm. Supernatants containing intracellular metabolites were lyophilized and stored at -80°C. These samples were reconstituted in 100 µl of 50% MeOH. LC-MS/MS analysis was carried out as described previously (71).

### Antibiotic minimum inhibitory concentration (MIC) measurements and synergy/checkerboard assays

MIC measurements by broth microdilutions were performed in accordance with CLSI methods for dilution susceptibility testing of staphylococci (72) with modifications as follows; guanosine, xanthosine or adenosine were supplemented into culture media at a final concentration of 200 µg/ml. Strains were first grown at 37°C on MHA 2% NaCl for 24 h and 5–10 colonies were resuspended in 0.85% saline before being adjusted to 0.5 McFarland standard (OD_600_=0.1). The cell suspensions were then diluted 1:20 in PBS and 10 µl used to inoculate 100 µl media (MHB 2% NaCl/ MHB 2% NaCl with 200 µg/ml guanosine) containing serially diluted antibiotics (oxacillin, 5-fluorouracil, 5-fluorouridine, sulfamethoxazole, trimethoprim) as indicated. The plates were incubated at 37 °C for 24 h and MIC values were recorded as the lowest antibiotic concentration where no growth was observed. Checkerboard/synergy assays performed as previously described (1), utilised 96 well plates in which one antibiotic/adjuvant was serially diluted vertically and the second antibiotic was serially diluted horizontally.

### Disc Diffusion synergy assays

Disc diffusions were carried out according to CLSI guidelines with the following modifications (73). Strains were grown overnight in TSB, serially diluted 1:100 (in fresh TSB) and grown for 4 h 37°C at 200rpm. The cultures were then adjusted to OD_600_=0.5 and spread evenly across 4mm thick MHA plates. Discs impregnated with antimicrobial drugs were placed onto plates and allowed to grow for 20 h. To measure synergy, two discs were placed 15mm apart and grown for 20 h to determine if there was increased growth inhibition in the region between the disks on MHA or MHA Gua (200 μg/ml).

### Antibiotic killing assays

Antibiotic killing assays were performed according to CLSI guidelines (72), as previously described (1). Briefly overnight cultures grown in MHB 2% NaCl were diluted 1:100 and grown for 3 h before being inoculated into 25 ml of MHB NaCl with or without antibiotics as indicated in 250 ml flasks at a starting cell density of approximately 1×10^6^ CFU/ml. The flasks were incubated at 35°C with shaking at 200 rpm and CFUs were enumerated on TSA plates after 0, 2, 4, 6, 8, 12, and 24 h. Glutathione (10 mM final concentration) was added as indicated.

### Biofilm eradication assays

Biofilm eradication assays were performed as described previously (74). Briefly, biofilms were grown in 2 ml BHI inoculated into the wells of 24-well tissue culture coated plates at 37℃ for 24 h before being carefully washed twice with PBS. BHI supplemented with antibiotics as indicated was then added to the biofilm wells and the biofilms incubated for a further 24 h at 37℃. The biofilms were dispersed by scraping, serially diluted and CFUs enumerated on TSA plates. The antibiotic concentrations used in these experiments was equal to or less than the C_max_ in humans as follows: oxacillin 200 µg/ml (17), sulfamethoxazole 78 µg/ml, trimethoprim 4 µg/ml (20), 5-fluorouracil 68 µg/ml (18), 5-fluorouridine 200 µg/ml (21) and vancomycin 50 µg/ml (22). An upper *in vivo* toxicity limit for guanosine has not been reported (23), and 200 µg/ml was used in this study. Three biological replicates were performed for each condition.

### Confocal microscopy using FM4-64 and HADA

For imaging of membranes and newly synthesized peptidoglycan, JE2 overnight cultures were inoculated 1:100 into 5 ml MHB cultures or MHB supplemented with oxacillin (1 µg/ml), 5-fluorouracil (1 µg/ml) and/or guanosine (200 µg/ml) as indicated. These cultures were then grown for 3 h before being incubated with HADA (500 µM final concentration) in the dark for 5 mins. Next the cells were pelleted for 2 min at 14,000 × g and resuspended in PBS supplemented with + 2 µg/ml FM4-64 at a cell density of OD_600_=1 before being incubated for 20 mins at 37°C. The cells were then washed twice with 1 ml PBS before being resuspended in PBS (OD_600_=1) and 5 µl aliquots were spot inoculated onto a 1% PBS agarose pad. The stained bacteria were then imaged at 1000× magnification using an Olympus LS FLUOVIEW Fv3000 Confocal Laser Scanning Microscope. Images were analysed using Fiji (ImageJ) software and representative images are shown.

### Antibiotic susceptibility spot dilution assays

Overnight cultures were adjusted to OD_600_ of 1 in PBS, and 10-fold serial dilutions made in a 96-well plate (20 µl of more concentrated suspension into 180 µl PBS). 5 µl of each dilution was spot inoculated onto MHA and MHA containing antibiotics +/- guanosine (200 µg/ml) +/- thymidine (50 μg/ml).

### Membrane potential measurements

The membrane potential of JE2 grown under different conditions was measured using 3,3′-diethyloxacarbocyanine iodide (DiOC2(3)) dye using flow cytometry using a previously described method (39, 41) with modifications. JE2 overnight cultures were inoculated 1:100 into 5 ml MHB or MHB supplemented with oxacillin (1 µg/ml), 5-fluorouracil (1 μg/ml) and/or guanosine (200 µg/ml) as indicated. These cultures were grown to OD_600_=0.8 before 1 ml aliquots were harvested by centrifugation at 14,000 rpm, and the cells washed twice with 1 ml of PBS. Thereafter, the cells were diluted to OD_600_=0.2 in PBS and a 100 µl of cell suspension aliquot added to 890 µl PBS and DiOC2(3) dye (30 µM final concentration). Carbonyl cyanide m-chlorophenylhydrazone (CCCP) (15 µM final concentration) was used as a control to collapse the membrane potential. 10,000 gated events were recorded per sample using BD Accuri™ C6 Plus Sampler Flow cytometer and BD Accuri™ C6 Plus Software, using the FL1 (green) and FL3 (red) channels. These experiments were performed on four biological replicates and the averages and standard deviation were plotted using FlowJo software V10.1, as described previously (39).

### ROS quantification

Intracellular reactive oxygen species (ROS) were quantified using H_2_DCFDA, this probe is taken up by living cells and oxidized by intracellular ROS producing a fluorescent readout (37). JE2 overnight cultures were inoculated 1:100 into 5 ml MHB or MHB supplemented with oxacillin (1 µg/ml), 5-fluorouracil (1 μg/ml) and/or guanosine (200 μg/ml) as indicated. These cultures were grown to OD_600_=0.8 and the cell density adjusted to OD_600_=0.2 in 1 ml volume before being stained with H_2_DCFDA fluorescent dye (final concentration 10 µM) as described previously (75). The cell suspensions were incubated for 10 mins, before 800 µl PBS supplemented with 100 mM EDTA was added to 200 µl of stained cells. This suspension was dispensed into flow tubes and wrapped in aluminium foil. ROS levels were measured by BD Accuri™ C6 Plus Sampler Flow cytometer and BD Accuri™ C6 Plus Software using the FL-1 channel, and 10,000 gated events (cells) were recorded. The data was analysed using FlowJo software V10.1.

## Author Contributions

Aaron Nolan: Conceptualization, Formal analysis, Funding acquisition, Investigation, Methodology, Writing - original draft, Writing - review & editing

Jessica B. Kelly: Formal analysis, Investigation, Methodology, Writing - review & editing

Jongsam Ahn: Formal analysis, Investigation, Methodology, Writing - review & editing Dhananjay Shinde: Formal analysis, Investigation, Methodology, Writing - review & editing

Vinai C. Thomas: Funding acquisition, Supervision, Writing - review & editing

Merve S. Zeden: Conceptualization, Formal analysis, Funding acquisition, Project administration, Investigation, Data curation, Methodology, Supervision, Writing - original draft, Writing - review & editing

James P. O’Gara: Conceptualization, Formal analysis, Funding acquisition, Project administration, Supervision, Writing - original draft, Writing - review & editing, Project management.

## Acknowledgements.

A.C.N. and J.P.O’G. were supported by grants from Research Ireland (19/FFP/6441) to J.P.O’G. and Thomas Crawford Hayes research award (2023) to A.C.N. J.B.K. and M.S.Z. were supported by Research Ireland (SFI-IRC) Pathway Fellowship 22/PATH-S/10804 and SDG Research Seed Funding from the University of Galway College of Science and Engineering to M.S.Z. J.A., D.S. and V.C.T. were funded by NIH/NIAID grants R01AI125588 and P01AI083211 (Metabolomics Core) to V.C.T. We are grateful to Dr Peter Owens from the Centre for Microscopy & Imaging facility and Dr Shirley Hanley from the Flow Cytometry Core facility in the Technology Services Directorate at the University of Galway for their technical support and assistance with confocal microscopy and flow cytometry, respectively. Mass spectrometry analyses were performed by the University of Nebraska Medical Center Multiomics Mass Spectrometry Core Facility (RRID: SCR_012539). The funders had no role in study design, data collection and interpretation, or the decision to submit the work for publication.

## Notes

### Competing Interest Statement

The authors have declared no competing interest.

## References

1. Nolan AC, Zeden MS, Kviatkovski I, Campbell C, Urwin L, Corrigan RM, Grundling A, O’Gara JP. 2023. Purine Nucleosides Interfere with c-di-AMP Levels and Act as Adjuvants To Re-Sensitize MRSA To beta-Lactam Antibiotics. mBio 14:e0247822.

2. Besier S, Ludwig A, Ohlsen K, Brade V, Wichelhaus TA. 2007. Molecular analysis of the thymidine-auxotrophic small colony variant phenotype of Staphylococcus aureus. Int J Med Microbiol 297:217–225.

3. Besier S, Zander J, Kahl BC, Kraiczy P, Brade V, Wichelhaus TA. 2008. The thymidine-dependent small-colony-variant phenotype is associated with hypermutability and antibiotic resistance in clinical Staphylococcus aureus isolates. Antimicrob Agents Chemother 52:2183–2189.

4. Chatterjee I, Kriegeskorte A, Fischer A, Deiwick S, Theimann N, Proctor RA, Peters G, Herrmann M, Kahl BC. 2008. In vivo mutations of thymidylate synthase (encoded by thyA) are responsible for thymidine dependency in clinical small-colony variants of Staphylococcus aureus. J Bacteriol 190:834–842.

5. Campbell ML, Marchaim D, Pogue JM, Sunkara B, Bheemreddy S, Bathina P, Pulluru H, Chugh N, Wilson MN, Moshos J, Ku K, Hayakawa K, Martin ET, Lephart PR, Rybak MJ, Kaye KS. 2012. Treatment of methicillin-resistant Staphylococcus aureus infections with a minimal inhibitory concentration of 2 mug/mL to vancomycin: old (trimethoprim/sulfamethoxazole) versus new (daptomycin or linezolid) agents. Ann Pharmacother 46:1587–1597.

6. Goldberg E, Bishara J. 2012. Contemporary unconventional clinical use of co-trimoxazole. Clin Microbiol Infect 18:8–17.

7. Kriegeskorte A, Block D, Drescher M, Windmuller N, Mellmann A, Baum C, Neumann C, Lore NI, Bragonzi A, Liebau E, Hertel P, Seggewiss J, Becker K, Proctor RA, Peters G, Kahl BC. 2014. Inactivation of thyA in Staphylococcus aureus attenuates virulence and has a strong impact on metabolism and virulence gene expression. mBio 5:e01447–01414.

8. Kriegeskorte A, Lore NI, Bragonzi A, Riva C, Kelkenberg M, Becker K, Proctor RA, Peters G, Kahl BC. 2015. Thymidine-Dependent Staphylococcus aureus Small-Colony Variants Are Induced by Trimethoprim-Sulfamethoxazole (SXT) and Have Increased Fitness during SXT Challenge. Antimicrob Agents Chemother 59:7265–7272.

9. Potter AD, Butrico CE, Ford CA, Curry JM, Trenary IA, Tummarakota SS, Hendrix AS, Young JD, Cassat JE. 2020. Host nutrient milieu drives an essential role for aspartate biosynthesis during invasive Staphylococcus aureus infection. Proc Natl Acad Sci U S A 117:12394–12401.

10. Cui L, Murakami H, Kuwahara-Arai K, Hanaki H, Hiramatsu K. 2000. Contribution of a thickened cell wall and its glutamine nonamidated component to the vancomycin resistance expressed by Staphylococcus aureus Mu50. Antimicrob Agents Chemother 44:2276–2285.

11. Buvelot H, Roth M, Jaquet V, Lozkhin A, Renzoni A, Bonetti EJ, Gaia N, Laumay F, Mollin M, Stasia MJ, Schrenzel J, Francois P, Krause KH. 2021. Hydrogen Peroxide Affects Growth of S. aureus Through Downregulation of Genes Involved in Pyrimidine Biosynthesis. Front Immunol 12:673985.

12. DiMaggio DA, Jr., Yeo WS, Brinsmade SR. 2025. Pyrimidine sufficiency is required for Sae two-component system signaling in Staphylococcus aureus. J Bacteriol 207:e0011525.

13. Connolly J, Boldock E, Prince LR, Renshaw SA, Whyte MK, Foster SJ. 2017. Identification of Staphylococcus aureus Factors Required for Pathogenicity and Growth in Human Blood. Infect Immun 85.

14. Patil M, Serhii K, Garzino F, Gobert Q, Giorgio S, Raimundo JM, Bolla JM, Camplo M. 2023. Synthesis and antimicrobial testing of 5-fluorouracil derivatives. Arch Pharm (Weinheim) 356:e2300103.

15. Hignett RC. 1964. The Incorporation of 5-Fluorouracil by Staphylococcus Aureus (Strain Duncan). Ii. Biochim Biophys Acta 91:584–591.

16. Singh V, Brecik M, Mukherjee R, Evans JC, Svetlikova Z, Blasko J, Surade S, Blackburn J, Warner DF, Mikusova K, Mizrahi V. 2015. The complex mechanism of antimycobacterial action of 5-fluorouracil. Chem Biol 22:63–75.

17. Neuville M, El-Helali N, Magalhaes E, Radjou A, Smonig R, Soubirou JF, Voiriot G, Le Monnier A, Ruckly S, Bouadma L, Sonneville R, Timsit JF, Mourvillier B. 2017. Systematic overdosing of oxa- and cloxacillin in severe infections treated in ICU: risk factors and side effects. Ann Intensive Care 7:34.

18. Casale F, Canaparo R, Serpe L, Muntoni E, Pepa CD, Costa M, Mairone L, Zara GP, Fornari G, Eandi M. 2004. Plasma concentrations of 5-fluorouracil and its metabolites in colon cancer patients. Pharmacol Res 50:173–179.

19. Nesseler N, Verdier MC, Launey Y, Malherbe A, Dermu M, Piau C, Flecher E, Tribut O, Malledant Y, Seguin P. 2014. High-dose continuous oxacillin infusion results in achievement of pharmacokinetics targets in critically ill patients with deep sternal wound infections following cardiac surgery. Antimicrob Agents Chemother 58:5448–5455.

20. Brown GR. 2014. Cotrimoxazole - optimal dosing in the critically ill. Ann Intensive Care 4:13.

21. Van Der Heyden SA, Highley MS, De Bruijn EA, Tjaden UR, Reeuwijk HJ, Van Slooten H, Van Oosterom AT, Maes RA. 1999. Pharmacokinetics and bioavailability of oral 5’-deoxy-5-fluorouridine in cancer patients. Br J Clin Pharmacol 47:351–356.

22. Barcia-Macay M, Lemaire S, Mingeot-Leclercq MP, Tulkens PM, Van Bambeke F. 2006. Evaluation of the extracellular and intracellular activities (human THP-1 macrophages) of telavancin versus vancomycin against methicillin-susceptible, methicillin-resistant, vancomycin-intermediate and vancomycin-resistant Staphylococcus aureus. J Antimicrob Chemother 58:1177–1184.

23. Schmidt AP, Bohmer AE, Schallenberger C, Antunes C, Tavares RG, Wofchuk ST, Elisabetsky E, Souza DO. 2010. Mechanisms involved in the antinociception induced by systemic administration of guanosine in mice. Br J Pharmacol 159:1247–1263.

24. Hawas S, Verderosa AD, Totsika M. 2022. Combination Therapies for Biofilm Inhibition and Eradication: A Comparative Review of Laboratory and Preclinical Studies. Front Cell Infect Microbiol 12:850030.

25. Brunke MS, Konrat K, Schaudinn C, Piening B, Pfeifer Y, Becker L, Schwebke I, Arvand M. 2022. Tolerance of biofilm of a carbapenem-resistant Klebsiella pneumoniae involved in a duodenoscopy-associated outbreak to the disinfectant used in reprocessing. Antimicrob Resist Infect Control 11:81.

26. Osland AM, Oastler C, Konrat K, Nesse LL, Brook E, Richter AM, Gosling RJ, Arvand M, Vestby LK. 2023. Evaluation of Disinfectant Efficacy against Biofilm-Residing Wild-Type Salmonella from the Porcine Industry. Antibiotics (Basel) 12.

27. Wang J, Han Y, Yang R, Zhao X. 2015. [Optimization of labeling and localizing bacterial membrane and nucleus with FM4-64 and Hoechst dyes]. Wei Sheng Wu Xue Bao 55:1068–1073.

28. Lund VA, Wacnik K, Turner RD, Cotterell BE, Walther CG, Fenn SJ, Grein F, Wollman AJ, Leake MC, Olivier N, Cadby A, Mesnage S, Jones S, Foster SJ. 2018. Molecular coordination of Staphylococcus aureus cell division. Elife 7.

29. Tinajero-Trejo M, Carnell O, Kabli AF, Pasquina-Lemonche L, Lafage L, Han A, Hobbs JK, Foster SJ. 2022. The Staphylococcus aureus cell division protein, DivIC, interacts with the cell wall and controls its biosynthesis. Commun Biol 5:1228.

30. Pinho MG, Errington J. 2003. Dispersed mode of Staphylococcus aureus cell wall synthesis in the absence of the division machinery. Mol Microbiol 50:871–881.

31. Lorian V. 1985. Low concentrations of antibiotics. J Antimicrob Chemother 15 Suppl A:15–26.

32. Thalso-Madsen I, Torrubia FR, Xu L, Petersen A, Jensen C, Frees D. 2019. The Sle1 Cell Wall Amidase Is Essential for beta-Lactam Resistance in Community-Acquired Methicillin-Resistant Staphylococcus aureus USA300. Antimicrob Agents Chemother 64.

33. Salamaga B, Kong L, Pasquina-Lemonche L, Lafage L, von Und Zur Muhlen M, Gibson JF, Grybchuk D, Tooke AK, Panchal V, Culp EJ, Tatham E, O’Kane ME, Catley TE, Renshaw SA, Wright GD, Plevka P, Bullough PA, Han A, Hobbs JK, Foster SJ. 2021. Demonstration of the role of cell wall homeostasis in Staphylococcus aureus growth and the action of bactericidal antibiotics. Proc Natl Acad Sci U S A 118.

34. Kim W, Zou G, Hari TPA, Wilt IK, Zhu W, Galle N, Faizi HA, Hendricks GL, Tori K, Pan W, Huang X, Steele AD, Csatary EE, Dekarske MM, Rosen JL, Ribeiro NQ, Lee K, Port J, Fuchs BB, Vlahovska PM, Wuest WM, Gao H, Ausubel FM, Mylonakis E. 2019. A selective membrane-targeting repurposed antibiotic with activity against persistent methicillin-resistant Staphylococcus aureus. Proc Natl Acad Sci U S A 116:16529–16534.

35. Hong Y, Li L, Luan G, Drlica K, Zhao X. 2017. Contribution of reactive oxygen species to thymineless death in Escherichia coli. Nat Microbiol 2:1667–1675.

36. Hong Y, Zeng J, Wang X, Drlica K, Zhao X. 2019. Post-stress bacterial cell death mediated by reactive oxygen species. Proc Natl Acad Sci U S A 116:10064–10071.

37. McBee ME, Chionh YH, Sharaf ML, Ho P, Cai MW, Dedon PC. 2017. Production of Superoxide in Bacteria Is Stress- and Cell State-Dependent: A Gating-Optimized Flow Cytometry Method that Minimizes ROS Measurement Artifacts with Fluorescent Dyes. Front Microbiol 8:459.

38. Painter KL, Strange E, Parkhill J, Bamford KB, Armstrong-James D, Edwards AM. 2015. Staphylococcus aureus adapts to oxidative stress by producing H2O2-resistant small-colony variants via the SOS response. Infect Immun 83:1830–1844.

39. Zeden MS, Schuster CF, Bowman L, Zhong Q, Williams HD, Gründling A. 2018. Cyclic di-adenosine monophosphate (c-di-AMP) is required for osmotic regulation in Staphylococcus aureus but dispensable for viability in anaerobic conditions. J Biol Chem 293:3180–3200.

40. Paez PL, Becerra MC, Albesa I. 2010. Effect of the association of reduced glutathione and ciprofloxacin on the antimicrobial activity in Staphylococcus aureus. FEMS Microbiol Lett 303:101–105.

41. Shapiro HM. 2008. Flow cytometry of bacterial membrane potential and permeability. Methods Mol Med 142:175–186.

42. Lu KY, Wagner NJ, Velez AZ, Ceppe A, Conlon BP, Muhlebach MS. 2023. Antibiotic Tolerance and Treatment Outcomes in Cystic Fibrosis Methicillin-Resistant Staphylococcus aureus Infections. Microbiol Spectr 11:e0406122.

43. Alborn WE, Jr., Allen NE, Preston DA. 1991. Daptomycin disrupts membrane potential in growing Staphylococcus aureus. Antimicrob Agents Chemother 35:2282–2287.

44. Labreche MJ, Lee GC, Attridge RT, Mortensen EM, Koeller J, Du LC, Nyren NR, Trevino LB, Trevino SB, Pena J, Mann MW, Munoz A, Marcos Y, Rocha G, Koretsky S, Esparza S, Finnie M, Dallas SD, Parchman ML, Frei CR. 2013. Treatment failure and costs in patients with methicillin-resistant Staphylococcus aureus (MRSA) skin and soft tissue infections: a South Texas Ambulatory Research Network (STARNet) study. J Am Board Fam Med 26:508–517.

45. Vicetti Miguel CP, Mejias A, Leber A, Sanchez PJ. 2019. A decade of antimicrobial resistance in Staphylococcus aureus: A single center experience. PLoS One 14:e0212029.

46. Proctor RA. 2008. Role of folate antagonists in the treatment of methicillin-resistant Staphylococcus aureus infection. Clin Infect Dis 46:584–593.

47. Khamash DF, Voskertchian A, Tamma PD, Akinboyo IC, Carroll KC, Milstone AM. 2019. Increasing Clindamycin and Trimethoprim-Sulfamethoxazole Resistance in Pediatric Staphylococcus aureus Infections. J Pediatric Infect Dis Soc 8:351–353.

48. Sato T, Ito R, Kawamura M, Fujimura S. 2022. The Risk of Emerging Resistance to Trimethoprim/Sulfamethoxazole in Staphylococcus aureus. Infect Drug Resist 15:4779–4784.

49. de Souza DC, Cogo LL, Dalla-Costa LM, Tomaz APO, Conte D, Riedi CA, Rosario Filho NA, Palmeiro JK. 2021. Emergence of Thymidine-Dependent Staphylococcus aureus Small-Colony Variants in Cystic Fibrosis Patients in Southern Brazil. Microbiol Spectr 9:e0061421.

50. El Yacoubi B, Bonnett S, Anderson JN, Swairjo MA, Iwata-Reuyl D, de Crecy-Lagard V. 2006. Discovery of a new prokaryotic type I GTP cyclohydrolase family. J Biol Chem 281:37586–37593.

51. Sankaran B, Bonnett SA, Shah K, Gabriel S, Reddy R, Schimmel P, Rodionov DA, de Crecy-Lagard V, Helmann JD, Iwata-Reuyl D, Swairjo MA. 2009. Zinc-independent folate biosynthesis: genetic, biochemical, and structural investigations reveal new metal dependence for GTP cyclohydrolase IB. J Bacteriol 191:6936–6949.

52. Rohrer S, Ehlert K, Tschierske M, Labischinski H, Berger-Bachi B. 1999. The essential Staphylococcus aureus gene fmhB is involved in the first step of peptidoglycan pentaglycine interpeptide formation. Proc Natl Acad Sci U S A 96:9351–9356.

53. Mikkelsen K, Sirisarn W, Alharbi O, Alharbi M, Liu H, Nohr-Meldgaard K, Mayer K, Vestergaard M, Gallagher LM, Derrick JP, McBain AJ, Biboy J, Vollmer W, O’Gara JP, Grunert T, Ingmer H, Xia G. 2021. The Novel Membrane-Associated Auxiliary Factors AuxA and AuxB Modulate beta-lactam Resistance in MRSA by stabilizing Lipoteichoic Acids. Int J Antimicrob Agents doi:10.1016/j.ijantimicag.2021.106283:106283.

54. Monteiro JM, Covas G, Rausch D, Filipe SR, Schneider T, Sahl HG, Pinho MG. 2019. The pentaglycine bridges of Staphylococcus aureus peptidoglycan are essential for cell integrity. Sci Rep 9:5010.

55. Longley DB, Harkin DP, Johnston PG. 2003. 5-fluorouracil: mechanisms of action and clinical strategies. Nat Rev Cancer 3:330–338.

56. Lennartz FE, Schwartbeck B, Dubbers A, Grosse-Onnebrink J, Kessler C, Kuster P, Schultingkemper H, Peters G, Kahl BC. 2019. The prevalence of Staphylococcus aureus with mucoid phenotype in the airways of patients with cystic fibrosis-A prospective study. Int J Med Microbiol 309:283–287.

57. Kovach K, Davis-Fields M, Irie Y, Jain K, Doorwar S, Vuong K, Dhamani N, Mohanty K, Touhami A, Gordon VD. 2017. Evolutionary adaptations of biofilms infecting cystic fibrosis lungs promote mechanical toughness by adjusting polysaccharide production. NPJ Biofilms Microbiomes 3:1.

58. Sedlmayer F, Woischnig AK, Unterreiner V, Fuchs F, Baeschlin D, Khanna N, Fussenegger M. 2021. 5-Fluorouracil blocks quorum-sensing of biofilm-embedded methicillin-resistant Staphylococcus aureus in mice. Nucleic Acids Res 49:e73.

59. McLeod JR, Harvey PA, Detweiler CS. 2021. An Oral Fluorouracil Prodrug, Capecitabine, Mitigates a Gram-Positive Systemic Infection in Mice. Microbiol Spectr 9:e0027521.

60. Twelves C, Wong A, Nowacki MP, Abt M, Burris H, 3rd, Carrato A, Cassidy J, Cervantes A, Fagerberg J, Georgoulias V, Husseini F, Jodrell D, Koralewski P, Kroning H, Maroun J, Marschner N, McKendrick J, Pawlicki M, Rosso R, Schuller J, Seitz JF, Stabuc B, Tujakowski J, Van Hazel G, Zaluski J, Scheithauer W. 2005. Capecitabine as adjuvant treatment for stage III colon cancer. N Engl J Med 352:2696–2704.

61. Younis W, Thangamani S, Seleem MN. 2015. Repurposing Non-Antimicrobial Drugs and Clinical Molecules to Treat Bacterial Infections. Curr Pharm Des 21:4106–4111.

62. Yeo WS, Arya R, Kim KK, Jeong H, Cho KH, Bae T. 2018. The FDA-approved anti-cancer drugs, streptozotocin and floxuridine, reduce the virulence of Staphylococcus aureus. Sci Rep 8:2521.

63. Oe C, Hayashi H, Hirata K, Kawaji K, Hashima F, Sasano M, Furuichi M, Usui E, Katsumi M, Suzuki Y, Nakajima C, Kaku M, Kodama EN. 2020. Pyrimidine Analogues as a New Class of Gram-Positive Antibiotics, Mainly Targeting Thymineless-Death Related Proteins. ACS Infect Dis 6:1490–1500.

64. Lu CH, Shiau CW, Chang YC, Kung HN, Wu JC, Lim CH, Yeo HH, Chang HC, Chien HS, Huang SH, Hung WK, Wei JR, Chiu HC. 2021. SC5005 dissipates the membrane potential to kill Staphylococcus aureus persisters without detectable resistance. J Antimicrob Chemother 76:2049–2056.

65. Saiman L, O’Keefe M, Graham PL, 3rd, Wu F, Said-Salim B, Kreiswirth B, LaSala A, Schlievert PM, Della-Latta P. 2003. Hospital transmission of community-acquired methicillin-resistant Staphylococcus aureus among postpartum women. Clin Infect Dis 37:1313–1319.

66. Gill SR, Fouts DE, Archer GL, Mongodin EF, Deboy RT, Ravel J, Paulsen IT, Kolonay JF, Brinkac L, Beanan M, Dodson RJ, Daugherty SC, Madupu R, Angiuoli SV, Durkin AS, Haft DH, Vamathevan J, Khouri H, Utterback T, Lee C, Dimitrov G, Jiang L, Qin H, Weidman J, Tran K, Kang K, Hance IR, Nelson KE, Fraser CM. 2005. Insights on evolution of virulence and resistance from the complete genome analysis of an early methicillin-resistant Staphylococcus aureus strain and a biofilm-producing methicillin-resistant Staphylococcus epidermidis strain. J Bacteriol 187:2426–2438.

67. O’Neill E, Pozzi C, Houston P, Smyth D, Humphreys H, Robinson DA, O’Gara JP. 2007. Association between methicillin susceptibility and biofilm regulation in Staphylococcus aureus isolates from device-related infections. J Clin Microbiol 45:1379–1388.

68. Fey PD, Endres JL, Yajjala VK, Widhelm TJ, Boissy RJ, Bose JL, Bayles KW. 2013. A genetic resource for rapid and comprehensive phenotype screening of nonessential Staphylococcus aureus genes. MBio 4:e00537–00512.

69. Liebeke M, Dorries K, Meyer H, Lalk M. 2012. Metabolome analysis of gram-positive bacteria such as Staphylococcus aureus by GC-MS and LC-MS. Methods Mol Biol 815:377–398.

70. Zeden MS, Gallagher LA, Bueno E, Nolan AC, Ahn J, Shinde D, Razvi F, Sladek M, Burke O, O’Neill E, Fey PD, Cava F, Thomas VC, O’Gara JP. 2023. Metabolic reprogramming and altered cell envelope characteristics in a pentose phosphate pathway mutant increases MRSA resistance to beta-lactam antibiotics. PLoS Pathog 19:e1011536.

71. Bulock LL, Ahn J, Shinde D, Pandey S, Sarmiento C, Thomas VC, Guda C, Bayles KW, Sadykov MR. 2022. Interplay of CodY and CcpA in Regulating Central Metabolism and Biofilm Formation in Staphylococcus aureus. J Bacteriol 204:e0061721.

72. CLSI. 2020. Performance Standards for Antimicrobial Susceptibility Testing. 30th ed. CLSI supplement M100., Wayne, PA.

73. CLSI. 2018. Performance Standards for Antimicrobial Disk Susceptibility Tests; approved standard—12th ed. M02-A13., Wayne, PA.

74. Durham PG, Sidders AE, Beam JE, Kedziora KM, Dayton PA, Conlon BP, Papadopoulou V, Rowe SE. 2021. Harnessing ultrasound-stimulated phase change contrast agents to improve antibiotic efficacy against methicillin-resistant Staphylococcus aureus biofilms. Biofilm 3:100049.

75. Podkowik M, Perault AI, Putzel G, Pountain A, Kim J, DuMont AL, Zwack EE, Ulrich RJ, Karagounis TK, Zhou C, Haag AF, Shenderovich J, Wasserman GA, Kwon J, Chen J, Richardson AR, Weiser JN, Nowosad CR, Lun DS, Parker D, Pironti A, Zhao X, Drlica K, Yanai I, Torres VJ, Shopsin B. 2024. Quorum-sensing agr system of Staphylococcus aureus primes gene expression for protection from lethal oxidative stress. Elife 12.

